# Systematic evaluation of cell-type deconvolution pipelines for sequencing-based bulk DNA methylomes

**DOI:** 10.1101/2021.11.29.470374

**Authors:** Yunhee Jeong, Lisa Barros de Andrade e Sousa, Dominik Thalmeier, Reka Toth, Marlene Ganslmeier, Kersten Breuer, Christoph Plass, Pavlo Lutsik

## Abstract

DNA methylation analysis by sequencing is becoming increasingly popular, yielding methylomes at single-base pair resolution. It has tremendous potential for cell-type heterogeneity analysis with intrinsic read-level information. Although diverse deconvolution methods were developed to infer cell-type composition based on bulk sequencing-based methylomes, the systematic evaluation has not been performed yet. Here, we thoroughly benchmark six previously published methods: Bayesian epiallele detection (BED), DXM, PRISM, csmFinder+coMethy, ClubCpG and MethylPurify, together with two array-based methods, MeDeCom and Houseman, as a comparison group. Sequencing-based deconvolution methods consist of two main steps, informative region selection and cell-type composition estimation, thus each was individually assessed. With these sophisticated evaluation, we demonstrate the method achieving the highest performance in different types of samples. We found that cell-type deconvolution performance is influenced by different factors depending on the number of cell types within the mixture. Finally, we propose a best-practice deconvolution strategy for sequencing data and limitations which need to be handled.

## 1 Introduction

Although single-cell analyses have been spearheading the progress in biomedicine lately, profiling of bulk samples is still in high demand for the analyses which cannot be readily accomplished using single-cell methods due to technical or cost-related reasons. For instance, single-cell methods are still too laborious and costly [1] for profiling of long-time biobanked samples or large tumor patient cohorts [2, 3, 4]. The bulk analysis of epigenetic modifications can demonstrate epigenetic variations in large cohorts and relate those with genomic and phenotypic characteristics [5, 6].

DNA methylation, particularly occurring at cytosines in CpG context in mammals, carries highly distinguishable cell-type-specific signals [7, 6]. The inheritability over cell divisions and the chemical stability make it significantly more accessible for profiling. These features of DNA methylation motivated research associating diseases with cell-type-specific DNA methylation signals. Such disease-associated methylation differences are identified as differentially methylated regions (DMRs). To this end, it was shown that tumor subtypes can be successfully classified based on bulk DNA methylation patterns of patient biopsies [8, 9, 10]. Nevertheless, cell-type-specific signals of DNA methylation can suffer from confounding factors, because DNA methylation is also associated with gender, age, various environmental influences etc. [11, 12]. These confounding factors make cell subpopulation analysis difficult by increasing the complexity of overall methylation patterns and obscuring cell-type-specific signals.

Even though there are *in vitro* techniques to purify individual cell types from bulk samples, such as cell sorting or cell enrichment, other spurious variation can be introduced into the samples during the experimental process and eventually further confound cell-type-specific methylation signals. As an alternative, cell-type deconvolution, a computational approach to infer the cell-type composition of bulk samples, is applicable for cell subpopulation analysis post-experimentally. Cell-type deconvolution has been extensively utilized to dissect array-based DNA methylation bulk data, e.g. Infinium 450K/EPIC microarrays [13, 14], that generates matrices of average methylation levels at specific loci in multiple samples (we refer to them as “array-shaped data”). *Reference-based* deconvolution methods infer cell-type proportions based on reference methylomes of purified cell populations. Various statistical methods and algorithms were employed to establish and fit reference-based deconvolution models including different forms of regressions [15, 16, 17], Expectation-Maximisation (EM) algorithm and deep neural networks [18, 19]. In contrast, *reference-free* deconvolution methods do not require reference data to infer the proportions of underlying cell types from methylomes. Numerous reference-free methods have been proposed by us and others [20, 21, 22, 23, 24], also broadly reviewed [25] and compared elsewhere [26]. Overall, reference-based and reference-free deconvolution of bulk DNA methylomes are well established and have been used routinely, in particular for the analysis of cellular composition in complex tumors [27, 28, 29, 30, 16].

More recently, DNA methylation sequencing approaches, such as reduced representation bisulfite sequencing (RRBS) [31] or whole genome bisulfite sequencing (WGBS) [32], have become increasingly popular owing to dropping sequencing costs, providing much broader genome coverage [33, 34, 35] and single-molecule resolution in the form of read-level methylation calls. Furthermore, single-cell bisulfite sequencing (scBS-seq) is being used increasingly [36, 37], particularly in conjunction with multi-omics analyses [38, 39]. However, few issues still have to be carefully managed in sequencing data. Since it has high genome coverage, the multiple methylation states from reads covering respective sites has to be refined and well-summarized. In addition, sequencing data consists of read-level methylomes that each CpG site includes multiple methylation state values from all reads covering the site. This abundance of information makes sequencing data analysis more intricate by concealing some cell-type-specific signals. Consequently, dealing with sequencing data highly depends on which genomic regions are analysed and how to interpret read-level information altogether.

Read-level DNA methylomes from sequencing data, in theory, should be more suitable and information-rich for cell composition inference in bulk samples, as compared to array-shaped data [28]. Diverse methods have been developed for cell-type deconvolution of sequencing-based methylome data, making use of the aforementioned advantages, foremost read-level resolution [40, 41, 42, 43, 44]. However, the methods published so far were evaluated based on different data sets and variable standards. Additionally, each method requires different preprocessing procedures and generates different forms of output which might confuse less experienced users. Nevertheless, previous studies mainly assessed array-based celltype deconvolution methods [26, 45]. With these distinctive advantages and rapidly growing demand of sequencing data analysis, a comprehensive, standardized and unbiased assessment of deconvolution methods targeting sequencing data has become more necessary.

In order to bridge this gap, we thoroughly compare and evaluate six previously published sequencing-based deconvolution algorithms: Bayesian epiallele detection (BED) [40], DXM [46], PRISM [42], MethylPurify [43], csmFinder + coMethy [41] and ClubCpG [44]. We evaluate their performance with respect to informative feature selection results and deconvolution accuracy under various experimental scenarios, using *in silico* mixtures of single-nucleus methylomes as well as realistic mixtures of tumor and normal WGBS data. Based on our analysis results, we finally propose efficient and trustworthy pipelines to deconvolve complex cell-mixture samples in different scenarios. To our knowledge, this benchmarking study is the first attempt to systematize and comprehensively evaluate the methodological developments in sequencing-based methylome deconvolution.

## 2 Results

### 2.1 Comparison of methodological designs

For our benchmarking study, we chose six published sequencing-based cell-type deconvolution methods (BED, DXM, PRISM, MethylPurify, csmFinder + coMethy and ClubCpG). As a comparison group, we added two cell-type deconvolution methods for array-shaped data (reference-based constrained projection method Houseman and our own reference-free method MeDeCom). To cope with the completely different input data formats, sequencing-based and array-based cell-type deconvolution procedures follow different pipelines in our project (**Figure 1**A).

**Figure 1:**
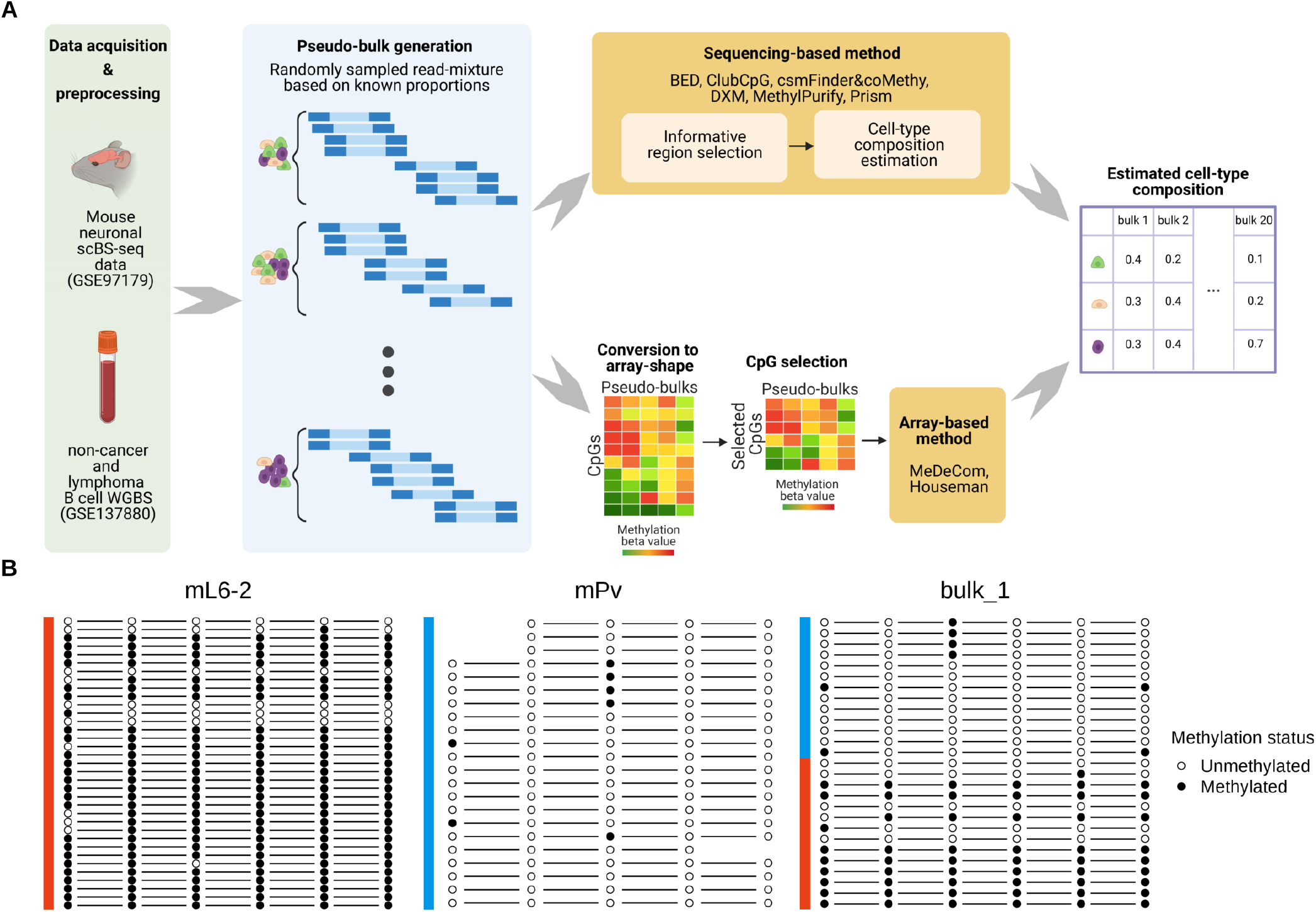
Characteristic overview of our cell-type deconvolution benchmarking. **A**. Overall scheme of cell-type deconvolution benchmarking. We synthesized *in silico* cell mixtures called pseudo-bulks by mixing up reads randomly sampled from mouse neuronal scBS-seq dataset and tumor WGBS dataset respectively. Sequencing-based cell-type deconvolution methods directly take sequencing data and select informative CpG sites (informative regions) supposedly related to cell-type heterogeneity (upper row pipeline). Then, cell-type proportions were estimated from selected CpGs. For array-based cell-type deconvolution methods, we converted sequencing data into array shape with pre-identified CpGs and the methods estimate cell-type compositions from the array (bottom row pipeline). **B**. Example of an informative region showing a cell-type-specific methylation pattern (chr1:75244319-75244379). Methylation pattern of reads (each row) overlapping with a specific region was extracted from three different samples, two pure cell-type samples (mL6-2 and mPv) and bulk_1 comprised of these cell types. Some missing methylation patterns in the figure occurred owing to the CpG sites not covered by each read. In this region, majority of reads from mL6-2 are fully methylated whereas most reads from mPv show a fully unmethylated pattern.

All compared sequencing-based methods consist of two common steps: informative region selection and cell-type composition estimation. In the informative region selection step, the sequencing-based celltype deconvolution methods filter out CpGs where the methylation patterns do not clearly demonstrate cell-type heterogeneity. Since sequencing data has significantly broader genome-wide coverage than array-shaped data, using all available CpG sites for deconvolution is not the most efficient strategy for cell-type composition estimation, associated with escalating computational complexity. In many genomic regions, methylation patterns are identical across all reads covering the CpG sites regardless of the cell type and thus non-informative. On the other hand, the some of patterns are over-complicated because of other confounding factors. Even, at such loci where read coverage is low, local cell-type population (distribution of reads in terms of cell type at the loci) usually does not correspond to global cell-type composition (cell-type composition in the entire bulk) due to the lack of reads. Informative region selection step alleviates these problems by clearing out these confounding methylation signals.

Initially, CpGs which are proximate to each other or overlap with a specific genomic region are grouped together. Here, we call each such group a *region*. After that, regions only satisfying particular criteria are selected for the next step and these selected regions are referred to as *informative regions* here. There are three common criteria considered in all our benchmarked methods: number of CpGs, region size and read coverage. The respective method retains only regions with an abundant number of overlapping CpGs, sufficient region length and high read coverage. The detailed informative region selection parameter values of each method are described in **Table 1**.

**Table 1:** Comparison of benchmarked deconvolution methods. Sequencing-based methods have three common criteria in the informative region selection: number of CpG, region size and read coverage. Some criteria were altered to be more suitable for dataset we used in our analyses. For array-based methods, we specifically designated CpG sites based on methylation variance or cell-type DMRs. Furthermore, we described differences in cell-type composition estimation step based on three criteria again: reference-requirements, number of detectable subpopulations and estimation scope.

| Method | Main data type | Informative region selection |  |  | Cell-type composition estimation |  |  |
| --- | --- | --- | --- | --- | --- | --- | --- |
|  |  | # CpG | Region Size (bp) | Coverage | Class | # Components | Estimation Scope |
| BED | RRBS | >4 | NA<sup>***</sup> | >20 | ref-based | 2 | local |
| ClubCpG | WGBS | >4* | 100 | >20* | ref-based | 2 or more | global |
| csmFinder + coMethy | WGBS | >4 | NA<sup>***</sup> | >10 | ref-free | 2 or more | global |
| DXM | Any kinds of BS-seq | Promoter and CpG island regions |  | >4 | ref-free | 2 or more | global |
| MethylPurify | WGBS | >10 | 300/200** | >10 | ref-free | 2 | local |
| Prism | RRBS | >4 | NA<sup>***</sup> | >20 | ref-free | 2 or more | local |
| Houseman | Methylation microarrays | CpGs overlapping with ctDMRs |  |  | ref-based | 2 or more | global |
| MeDeCom | Methylation microarrays | CpGs showing high methylation variance over covering reads |  |  | ref-free | 2 or more | global |

\* The original setup in [44] is 2 for minimum number of CpG and 10 for minimum read coverage.
\*\* The original setup in [43] is 300 bp for region size. In tumor-normal pseudo-bulk analysis, we changed region size parameter value to 200 bp.
\*\*\* BED, csmFinder and Prism do not require specific region size.

For the two array-based methods, however, informative region selection is not a required step, since DNA methylation array-shaped data is already designed as a matrix of CpG sites by samples. Therefore, we converted our sequencing data to an array shape with pre-designated CpG sites. We described CpG site selection procedure for each array-based method in **Methods**.

With the selected informative regions (sequencing-based methods) or pre-designated CpG sites (array-based methods), cell-type deconvolution methods predict cell-type proportions within given input cell-mixture samples. This step can be characterized by three main specifications: reference requirement, number of detectable cell types/subpopulations and estimation scope (**Table 1**).

#### Requirement of reference methylomes

Among the methods we benchmarked, BED, ClubCpG and Houseman are categorized as reference-based methods. BED and Houseman require methylome profiles of pure cell types. ClubCpG fits a regression model to training cell-mixture bulk data given together with known cell-type composition.

#### Number of detectable components

We also categorized the benchmarked methods with respect to whether the method specifically targets tumor samples or not. The difference between tumor targeting and broadly applicable methods is their assumption about cell-type composition within given samples, and the methods targeting tumor are generally referred as *tumor purity estimation methods*. These assume that only two subpopulations comprise given cell mixtures, tumor and healthy stroma, whereas normal cell-type deconvolution method does not limit the number of subpopulations. Among the methods we considered, BED and MethylPurify were developed particularly for the analysis of tumor samples. So, we additionally tested our benchmarked methods with realistically simulated tumor-normal bulk samples, since those might be better suited to deal with abnormal methylation patterns from tumor cell types.

#### Estimation scope

Exploring all the methods, we have found a prevalent computational approach which summarizes methylation patterns across all of selected informative regions and compute the final estimates. BED, MethylPurify and PRISM *locally* calculate statistics in respective informative regions and predict compositions from the peak of region-wise estimated cell-type composition distribution. On the other hand, ClubCpG, coMethy, MeDeCom and Houseman *globally* calculate final cell-type proportions directly over all selected informative regions.

### 2.2 Benchmarking study design

To estimate the influence of each step upon the final result, we comprehensively assess all methods not only with the final cell-type composition estimation results, but also with the interim results of selected informative regions for deconvolution.

Our benchmarking analysis was performed with three different datasets to test the capability of methods in various biological scenarios, two and five cell-type mouse neuronal pseudo-bulks and tumor pseudo-bulk (**Table 2**). Firstly, we used single-nucleus bisulfite sequencing data of mouse neuron population from Luo et. al [47] to generate *in silico* pseudo-bulk read/cell mixtures for two and five cell types. Secondly, to test methods in a realistic scenario of tumor bulk deconvolution, we created *in silico* mixtures of WGBS methylomes from normal non-cancer B-cell and B-cell lymphoma samples from Do et. al [48].

**Table 2:** Details of pseudo-bulk datasets. We created three datasets of pseudo-bulk samples for evaluating our benchmarked methods. Two datasets are comprised of two and five cell-types of mouse neuronal single-cell BS-seq data each, and one dataset was created with tumor and normal cell types. We generated 20 pseudo-bulk samples in each dataset.

|  | Resource | # bulks | Cell types | Biological sample | Sequencing protocol |
| --- | --- | --- | --- | --- | --- |
| 2 cell-type mouse neuronal pseudo-bulk | GSE97179<br>[47] | 20 | mL6-2, mPv | Mouse neuron | scBS-seq<br>(Illumina HiSeq 4000) |
| 5 cell-type mouse neuronal pseudo-bulk | GSE97179<br>[47] | 20 | mL6-2, mPv, mDL-2,<br>mL2-3, mL5-1 | Mouse neuron | scBS-seq<br>(Illumina HiSeq 4000) |
| Tumor-normal pseudo-bulk | GSE137880<br>[48] | 20 | Normal B-cell,<br>B-cell lymphoma | B-cell | WGBS<br>(Illumina NovaSeq 6000) |

In the analysis of five cell-type pseudo-bulk group, we excluded BED and MethylPurify, because tumor purity estimation methods are not eligible to detect more than two cell types. Although Prism is not a tumor purity estimation method, it failed in detecting five cell types so we also excluded it from five cell-type pseudo-bulk analysis.

### 2.3 Evaluation of the informative region selection methods using cell-type DMRs

We hypothesized that ideal informative region selection results should overlap with cell-type DMRs (ctDMRs), i.e. genomic regions showing significant methylation pattern differences between cell types (**Figure 1**B). Conversely, informative region selection results with low similarity to ctDMRs are unlikely to supply enough cell-type-specific signals. Hence, we assessed informative region selection results primarily by comparing them with ctDMRs. The ctDMRs were generate by comparing one pure cell-type bulk to all others (details in **Methods**).

For two cell-type neuronal pseudo-bulks, BED detected a significant number of informative regions overlapping with ctDMRs, but csmFinder detected the highest number of overlaps in tumor pseudo-bulks. (**Figure 2**A, B) In five cell-type mouse neuronal pseudo-bulk analysis, ClubCpG showed the highest number of overlaps with ctDMRs over all cell types (**Supplementary Figure 1**).

**Figure 2:**
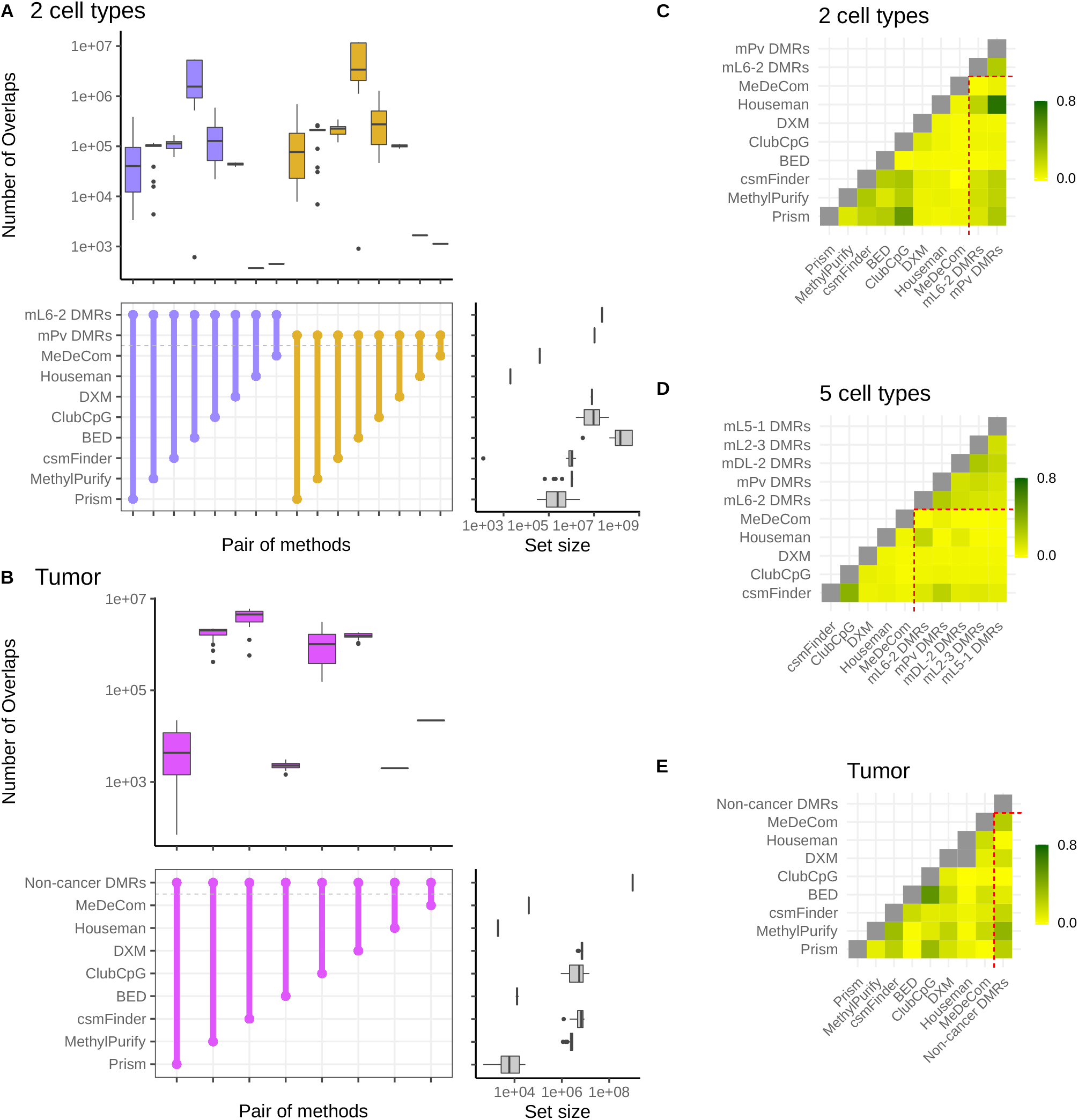
Plots for informative region selection. **A, B**. Overlaps between ctDMRs for **(A)** two cell-type mouse neuronal pseudo-bulks and **(B)** tumor pseudo-bulks. The colored box plots at the top of each graph show the number of overlaps between a pair of methods connected at the middle, across all pseudo-bulks, and the grey box plots at the right side display the number of informative regions detected by each method or the number of regions in ctDMRs. Overlap sizes were calculated with respect to the number of bases. In each plot, different ctDMRs are distinguished by different colors. **C-E**. Genomic correlation between ctDMRs and selected informative regions in **(C)** two cell-type mouse neuronal pseudo-bulks, **(D)** five cell-type mouse neuronal pseudo-bulks and **(E)** tumor pseudo-bulks. Higher genomic correlation means higher similarity between ctDMRs and selected informative regions with respect to the number of overlaps and the proximity.

Since the size of the selected informative region set is different among methods, the number of overlaps may not exactly correspond to how similar each selected informative region set is to ctDMRs. We found that the set size of selected informative regions and the number of overlaps with ctDMRs have a strong correlation in all datasets (**Supplementary Figure 4**). Consequently, our results clearly showed that selecting more informative regions possibly increases the number of overlaps with ctDMRs even though the selected region set may also include much more non-overlapping regions. Thus, we conducted another statistical analyses to compare the similarity between selected informative regions and ctDMRs.

Favorov et al. designed a statistic called *genomic correlation* which measures the distributions of distances between two sets of genomic regions [49] (details in the **Methods**). This statistic is [0,1] interval-bound, with lower value indicating larger distance between genomic regions from the two compared sets. We evaluated the informative region selection results based on the proximity to ctDMRs using genomic correlation score.

Although informative regions of BED overlapped the largest number of ctDMRs in two cell-type mouse neuronal pseudo-bulk samples, their genomic correlation with each ctDMR set was the lowest (**Figure 2**C). This implies that ctDMRs comprised a very small fraction of the informative regions selected by BED. In contrast, csmFinder, Prism and MethylPurify showed high genomic correlation with ctDMRs in both two cell-type mouse neuronal and tumor pseudo-bulks, meaning that ctDMRs take a large portion of the selected regions in spite of the small absolute total number. For five cell-type mouse neuronal pseudo-bulks, csmFinder yielded the highest genomic correlation with all ctDMRs compared to other sequencing-based methods, ClubCpG and DXM.

We further explored genome annotations of the selected informative regions (**Supplementary Figure 7-9**). Compared to ctDMRs, the selected informative regions included noticeably larger amount of promoter regions in the all types of pseudo-bulk analyses with the exception of BED in two cell-type mouse neuronal pseudo-bulk result. Considering the methylation located in a gene promoter is able to regulate different gene expression according to cell types, we suppose that the informative region selection step of the benchmarked methods well identifies methylation patterns contributing to deciding cell-type identity.

We also calculated the methylation level difference between two cell types particularly with bi-component (two cell-type mouse neuronal and tumor) pseudo-bulks. Within each set of selected informative regions, we extracted methylation beta value at CpGs from two pure cell-type methylomes and calculated the difference. The distribution of differences was calculated in each bulk (**Supplementary Figure 5 and 6**). If the selected regions covers CpGs with cell-type specific methylation patterns, the methylation level difference value must be close to either 1 or -1 depending on hypomethylated cell type at the CpG site. Cell-type DMRs, as expected, mostly covers CpGs with the absolute methylation level difference of 1. Regions designated for array-based methods also showed the methylation difference distribution peaking at -1 and 1, as ctDMRs and CpGs with the highest variance of methylation value were given to Houseman’s method and MeDeCom respectively. However, CpGs selected by sequencing-based methods dominantly presented the methylation level difference 0, especially for two cell-type mouse neuronal pseudo-bulks. From tumor pseudo-bulk samples, csmFinder, MethylPurify and Prism were more successful in detecting CpGs showing high methylation difference between non-cancer and B-cell lymphoma cell types.

In conclusion, we found that sequencing-based methods can detect ctDMRs through their informative region selection step. However, despite the large number of overlaps with ctDMRs, the selected informative region set still include some uninformative genomic regions, distant from ctDMRs or showing zero methylation level difference across different cell types. Methylation patterns from such regions may hinder accurate cell-type deconvolution, if it is not handled during the cell-type composition estimation step.

### 2.4 Mouse neuronal single-cell pseudo-bulk cell-type deconvolution

As explained above, we evaluated all methods with two groups of mouse neuronal single-cell pseudo-bulk samples. Absolute error value between the ground-truth and estimated value is a main score for the cell-type composition estimation step. Reference-based and reference-free methods were evaluated separately for a fair comparison in terms of additional prior information.

Our results showed that reference-based methods, with the exception of BED, perform better than reference-free methods in both groups (**Figure 3**A and B). Furthermore, we analysed predicted proportion of each cell type within each pseudo-bulk sample (**Figure 3**C and D). We realized that ClubCpG can produce an cell-type proportion estimate lower than 0 or higher than 1, when ground-truth proportion is relatively low or high. This is because ClubCpG does not restrict its prediction value between 0 and 1. Yet, other methods successfully made all predictions in the range of 0 and 1.

**Figure 3:**
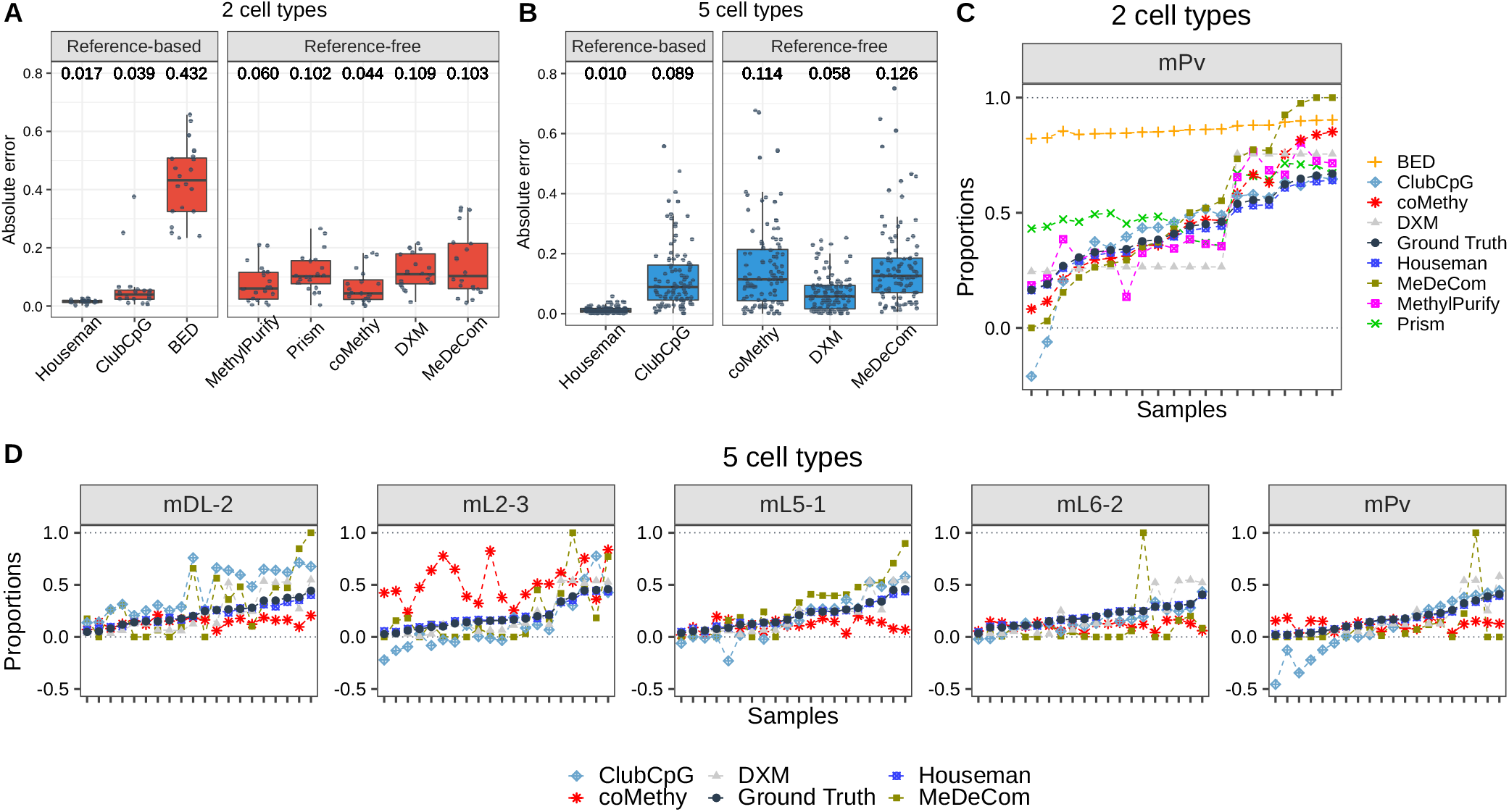
Cell-type composition estimation for mouse neuronal single-cell pseudo-bulk scenario. **A, B**. Absolute error between ground-truth and estimated cell-type proportion, calculated in each sample and each cell type, for **(A)** two cell-type and **(B)** five cell-type mouse neuronal pseudo-bulks. Black line at the middle refers to the median and the both ends of bar are the first and third quantiles. Numbers above box plots indicate the median value. **C, D**. Estimated cell-type proportions and ground-truth values (black line) ordered with respect to the ground-truth proportion, in **(C)** two cell-type and **(D)** five cell-type mouse neuronal pseudo-bulks. For two cell-type pseudo-bulks, only results of mPv cell type are shown to avoid duplicates of cell-type proportions in bi-component mixtures.

In the two cell-type pseudo-bulk analysis, the best accuracy was achieved by Houseman’s method requiring pure cell-type methylome profiles. Among reference-free methods, coMethy performed best with a median absolute error value of 0.044, even though MethylPurify and Prism inferred more accurate values for bulks with high proportion of mPv.

Among reference-based methods, the pseudo-bulk with five cell types showed the same results as with two cell types: the lowest median absolute error was recorded by Houseman. However, among reference-free methods, DXM performed best with the median absolute error value 0.058. In bulk-wise performances separated by cell types. mL2-3 was the the most difficult cell type to estimate for coMethy, but ClubCpG and MeDeCom recorded the lowest accuracy in mDL-2. Even though all other methods showed better performance with two cell-type pseudo-bulks, Houseman and DXM exhibited lower median absolute error with five cell-type pseudo-bulks than with two cell-type pseudo-bulks.

#### 2.4.1 Tumor-normal cell mixture deconvolution

Tumor heterogeneity analysis is one of the most crucial and broadly conducted research in cancer biology, but it is significantly harder due to the additional sources of variation. Reflecting the uniqueness of tumor tissues, BED and MethylPurify are particularly designed for tumor purity estimation of tumor-normal cell mixture as described in **Table 1**. For this reason, we additionally assessed the benchmarked methods using pseudo-bulk samples comprised of B-cell lymphoma and normal non-cancer B-cell (**Figure 4**). In order to cover the high variation in tumor, we used WGBS data derived from cell line with a pure cell population, for generating pseudo-bulks rather than sparse scBS-seq data.

**Figure 4:**
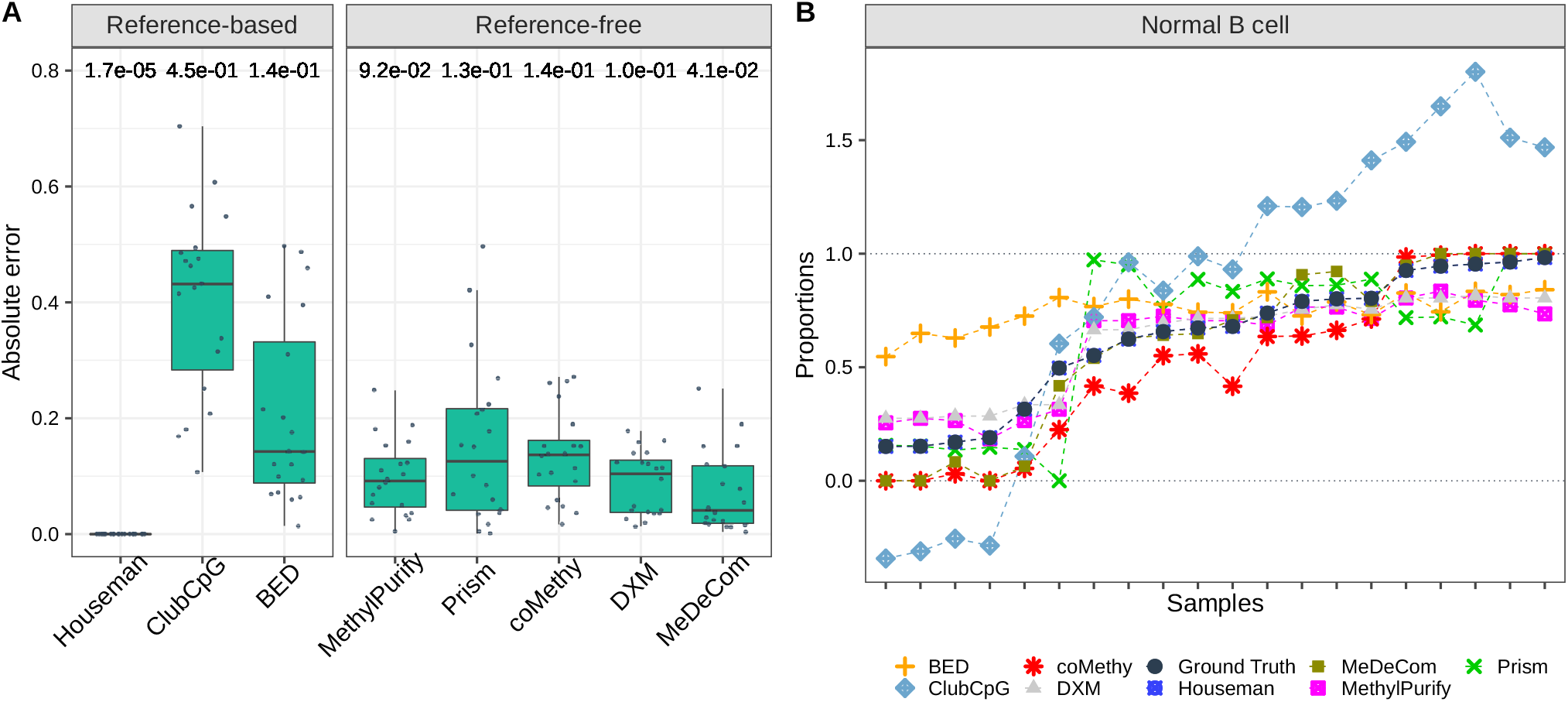
Cell-type composition estimation results of tumor pseudo-bulk scenario. **A**. Absolute error between estimated and ground-truth cell-type proportions. **B**. Estimated proportions by each method and ground-truth values in order. All details of two plots are the same as in Figure 3. Again, we present the results only for normal B-cell due to the symmetric cell-type proportions in bi-component mixtures.

In the reference-based setting, Houseman’s method again showed the best performance in tumor celltype deconvolution. Compared to mouse neuronal pseudo-bulk deconvolution, Houseman’s method accomplished the lowest median absolute error of 1.7*e*^−05^ in tumor cell-type deconvolution. Among reference-free methods, MeDeCom estimated tumor pseudo-bulk cell-type compositions with the lowest median absolute error. Sample-wise performance for one cell type also showed that Houseman estimated the most accurate proportions for all samples and ClubCpG exceeded the range of 0 and 1 for extreme proportions similarly to mouse neuronal pseudo-bulk analysis results.

### 2.5 Identifying factors that influence cell-type deconvolution performance

Based on the results so far, we investigated whether informative region selection results have detectable influence upon cell-type composition estimation. For this, We compared the rank of mean absolute error together with the rank of mean genomic correlation separately (**Figure 5**A and **Supplementary Figure 2**). Both absolute error and genomic correlation results from a method were averaged across all samples but separately in each cell type, whereafter we ranked the methods with the averaged values. After ranking the methods, we excluded array-based methods whose pipeline does not have a informative region selection. For tumor pseudo-bulks, two cell-type results from each method are assigned in a same rank of mean genomic correlation, since B-cell non-cancer and lymphoma cell types involve same ctDMRs. Consequently, in two cell-type mouse neuronal and tumor pseudo-bulk samples, the accuracy of cell-type composition estimation (complemented of the mean absolute error value) tends to be proportional to the performance of genomic correlation between selected informative regions and ctDMRs. From this result, even though all methods are designed with different algorithms, we claim that detecting CpGs overlapping with ctDMRs significantly contributes to the performance of cell-type deconvolution generally in the cell mixture of bi-components.

**Figure 5:**
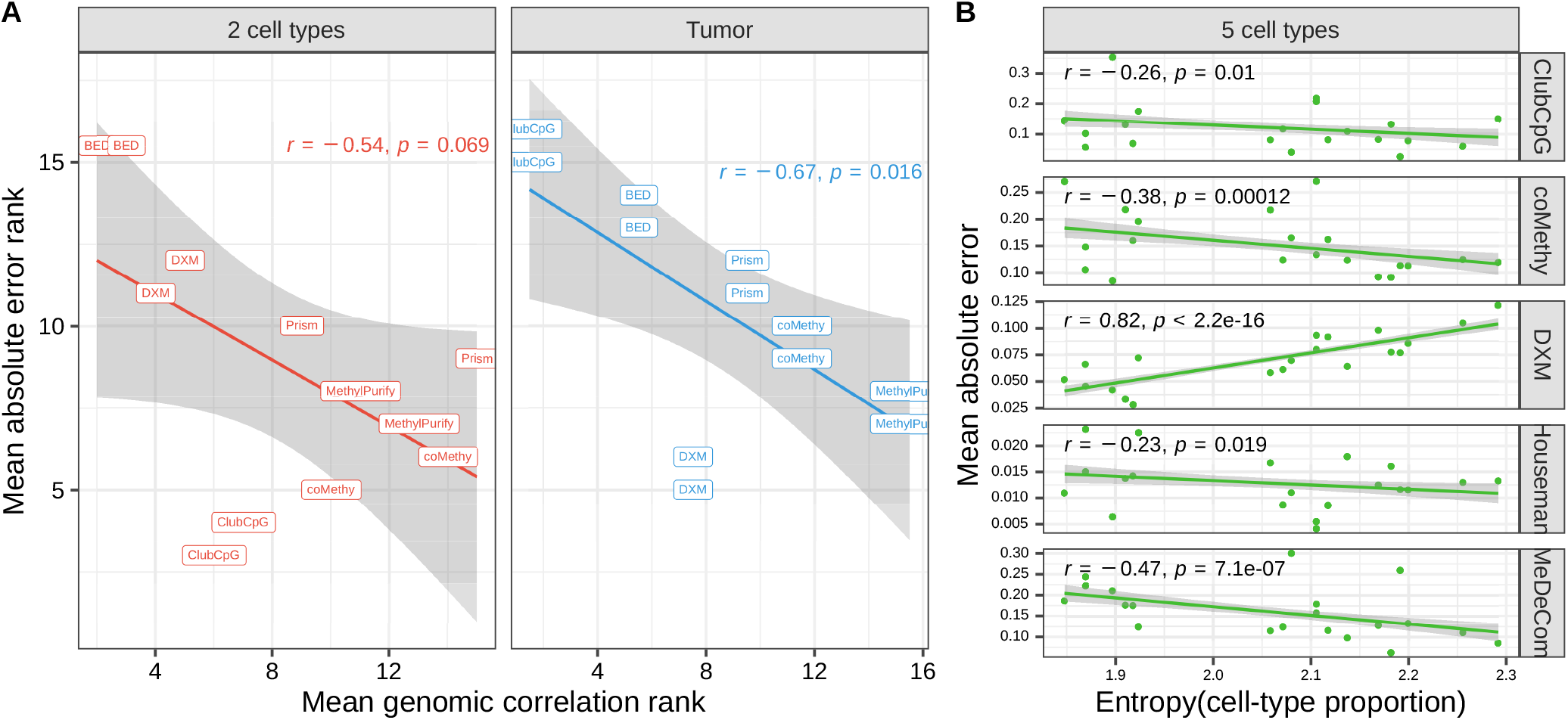
Influential factors in cell-type deconvolution performance. **A**. Mean absolute error vs mean genomic correlation between selected informative regions and ctDMRs. The points indicate results of each cell type from each deconvolution method. **B**. Mean absolute error vs entropy of cell-type proportions in each bulk sample. In both plots, we fitted dots in a linear function (the line with grey background).

<We additionally discovered that all methods except DXM could infer more accurate cell-type proportions in the mixtures with more balanced cell-type composition in five cell-type pseudo-bulks (**Figure 5**B). This was determined using the entropy value which measures the uniformity of given proportions or given distribution (details in **Methods**). The entropy of cell-type distribution was negatively correlated with the mean absolute error showing the p-value lower than 0.05 in the results of ClubCpG, coMethy, Houseman and MeDeCom. We presume that, in cell-type mixtures with low entropy values where cell population is extremely distributed, minor cell types may not provide enough cell-type specific signals. Also, extreme value of proportions in low entropy of cell-type distribution, very close to 0 or 1, could deteriorate the performance. Nevertheless, DXM showed the reversed results from the four methods. The lower entropy of pseudo-bulks are given, it showed the better performance. We suppose that the design of DXM algorithm searching the best fitting out of randomly generated distributions rather than gradually fitting a model becomes more powerful in the extreme distribution of cell types by disregarding regularization.

In both two cell-type deconvolution scenarios, not all methods performed better with higher entropy of samples. For instance, Houseman in the two cell-type pseudo-bulk analysis and coMethy, Prism in tumor pseudo-bulk analysis rather showed positive correlation between two values (**Supplementary Figure 3**). This might be caused by some bulks with high entropy, where cell types are more uniformly distributed, resulting in more complicated methylation patterns as similar amount of cell-type specific signals from each cell type.

## 3 Discussion

Here we extensively reviewed and assessed five sequencing-based cell-type deconvolution methods with standardized measurements for unbiased evaluation. Two more array-based deconvolution methods, MeDeCom and Houseman, were also added to the analyses as a comparison group, to evaluate the effectiveness of the benchmarked methods to leverage the unique properties of sequencing data.

In order to reflect various biological scenarios, our analyses were done with two different datasets, mouse neuronal scBS-seq dataset and B-cell tumor WGBS dataset. We generated pseudo-bulk samples by merging randomly sampled reads from pure cell-type samples in each dataset. For mouse neuronal pseudo-bulk samples, we generate mixtures of two and five cell types respectively to study scenario of different number of subpopulations.

All sequencing-based methods we evaluated include two essential steps. First, each method selects genomic regions which are considered to show clear cell-type-specific methylation patterns (informative region selection). In the second step, final cell-type composition is inferred based on the selected regions (cell-type composition estimation). Thus, we evaluated the performance separately in respective steps and finally examined the influential factors in cell-type deconvolution performance.

While evaluating informative region selection, we regarded ctDMRs as the gold-standard cell-type-specific loci and compared them with informative regions selected by each method. Although, in mouse neuronal pseudo-bulk analyses, ClubCpG detected the largest number of overlaps with ctDMRs, csmFinder showed the best genomic correlation. This is because that large number of overlaps can be caused by the large number of total selected informative regions, not by the capability of detecting cell-type-specific regions (**Supplementary Figure 1**).

To assess cell-type composition estimation results, we calculated the absolute error between the estimated and ground-truth proportion of each cell type in each sample. Since introducing prior knowledge from reference data ordinarily improves estimation performance, for the evaluation we grouped methods according to whether reference data is required or not. Among reference-based methods, Houseman’s method strongly outperformed other methods. In the comparison of reference-free methods, coMethy inferred the most accurate cell-type compositions in both two and five cell-type pseudo-bulks.

Cancer-associated DNA methylation patterns are particularly aberrant and the cell subpopultions in cancerous tissues containing both healthy normal and tumor cell types often bring more complicated composition [50]. Therefore, we separately evaluated cell-type deconvolution results in tumor pseudo-bulk samples generated as described above. In informative region selection analysis, csmFinder not only showed the large number of overlaps with the ctDMRs, but also reached the highest genomic correlation. For cell-type composition estimation, Houseman and MeDeCom outperformed all others in reference-based methods and in reference-free methods each.

Lastly, we have noticed the significance of selected informative regions for the overall cell-type deconvolution performance. As shown in **Figure 5**, mean absolute error and genomic correlation of evaluated methods have a negative rank correlation in bi-component pseudo-bulks. This result highlights that the more similar informative regions to ctDMRs a method can detect, the better performance the method would achieve in cell-type composition estimation for cell mixtures comprised of two distinct subpopulations. Yet, in five cell-type mouse neuronal pseudo-bulks, entropy of cell-type distribution showed a negative correlation with the absolute error except DXM (**Figure 5**). This implies that the distribution of subpopulations can be more influential in cell-type deconvolution of complex bulk samples.

Although our benchmarked methods carried out rational cell-type deconvolution results in most analyses, there are still some limitations which have to be resolved in sequencing-based cell-type deconvolution. Firstly, based on our results, sequencing-based methods did not outperform array-based methods in cell-type composition estimation. Not only Houseman’s method performed best of all reference-based methods, but also MeDeCom achieved the lowest median absolute error among reference-free methods in the tumor pseudo-bulk analysis. One of the factors making sequencing-based approach intricate is the high complexity of sequencing data itself. The complexity arises because, unlike array data consisting of summarized methylation beta value, sequenicng data involves a methylation state from multiple reads at each CpG site. Thus, sequencing-based method should be capable of eliminating redundant methylation patterns and statistically well-modeling subpopulation distribution with remaining informative methylation patterns.

With the leverage of single-cell profiling, it was convincingly demonstrated that the variable composition of numerous normal cell types and tumor cell subclones underlies intra-tumor heterogeneity [51, 52, 53]. However, tumor purity estimation methods in our analyses could only consider tumor samples as binary mixtures of tumor and normal cell subpopulations. These results are not able to explain the actual complexity of tumor microenvironment which has been widely investigated, particularly in relation to success of various therapeutic strategies, e.g. immunotherapy [54]. Therefore, to support the upcoming tumor studies with the analysis of cell-type composition variation, more advanced tumor-targeting cell-type deconvolution methods addressing multi-component intra-tumor heterogeneity need to be implemented.

Last but not least, the software availability, sustainability and deployment should be considered important when cell-type deconvolution tools are implemented and released. To this end, we found that some of our benchmarked methods have significant methodological and technical limitations. For example, PRISM and BED do not ensure its feasibility to other types of data than RRBS. Furthermore, most of benchmarked methods are available only for samples aligned with *Bismark* [55]. The limited availability, complexity of deployment and lack of input standardization eventually hinder efficient utilization of evaluated methods for real-life analyses. In addition, software maintenance is another crucial issue to develop a sustainable celltype deconvolution tool. Importantly, to be able to execute the tools, we had to implement multiple bugfixes. Given that bisulfite sequencing technology has been rapidly innovated recently [33, 36], sequencing-based cell-type deconvolution tools should be well maintained and updated to keep the usefulness correspondingly.

Taken together, our analysis results suggest a clear paradigm of how to conduct cell-type deconvolution for sequencing data. It will pave the way towards more accurate cell-type composition inference and more precise analysis of cell-type-specific methylation patterns to allow further method development and improvement. Because of the intrinsic benefit of read-level information, which provides detailed methylation states at each CpG, it should be possible to accomplish better accuracy in the inference of cell-type composition from sequencing-based DNA methylation data compared to summarized (arrayshaped) data, something which currently available tools cannot achieve yet. For the future work, it has to be more thoroughly designed how to extract cell-type-specific signals and adjust for confounding factors affecting sequencing data. Precise cell-type deconvolution of sequencing data can broaden the range of available analyses and support a strong cell population examination tool in diversified clinical and biomedical applications.

## 4 Methods

### 4.1 Benchmarking datasets

In order to evaluate the sequencing-based deconvolution methods, we generated synthetic cell-mixture samples called pseudo-bulk. For the pseudo-bulk generation, we have chosen two different datasets, mouse brain single-cell methylC-seq data [47] and B-cell tumor WGBS data [48].

#### 4.1.1 Single-cell methylome data processing

3377 single-nucleus methylomes derived from 8-week old mouse cortex tissue were downloaded from the Gene Expressiong Omnibus (GEO) with the accession number GSE97179. This dataset was created through high-throughput single-nucleus methylome sequencing (snmC-seq). We firstly trimmed reads twice to remove sequencing adaptors, random primer index sequence and C/T tail attached, using Adaptase with Cutadapt 2.6 [56]. Trimmed reads were aligned to the *mm10* reference genome using Bismark 0.22.3 [55]. After the alignment, we sorted the result BAM files using samtools 1.9 [57] and removed duplicated reads using picard MarkDuplicates 1.141. Finally, the reads whose mapping quality is lower than 30 were filtered out with samtools 1.9 again. Parameters used in each step and details about these processes are clarified in **Supplementary Table 1**.

#### 4.1.2 Tumor WGBS data processing

To generate realistic tumor-normal cell mixtures, we downloaded diffuse large B-cell lymphoma and normal non-cancer B-cell WGBS data from one subject each (GEO accession number GSE137880) [58]. Trimming was conducted using Trim Galore 0.6.6, then reads were aligned by Bismark 0.22.3 [55] in paired-end mode. Only reads not aligned in paired-end mode were re-aligned in single-end mode, and all reads were merged and sorted through samtools 1.9 [57]. At the end, duplicated reads were removed by picard Mark Dubplicates 1.141. Detailed pipeline is clarified in **Supplementary Table 1**.

#### 4.1.3 Pseudo-bulk generation

We created pseudo-bulks by merging reads randomly sampled from the aligned mouse brain single-nucleus methylome data and B-cell WGBS data. The cell-type proportion for bulks were decided based on *Dirichlet distribution* which can mimic diverse biological scenarios. For mouse neuron pseudo-bulk, five cell types including both excitatory (mDL-2, mL2-3, mL5-1 and mL6-2) and inhibitory (mPv) neuron classes were chosen. These cell types are distinctly clustered in the two-dimensional t-SNE visualization calculated by Luo et. al. [47] and secure sufficient single-cell samples. For tumor pseudo-bulk, diffuse large B-cell lymphoma and normal B-cell cell-types were mixed. We used *generateExample* function from R package MeDeCom (https://rdrr.io/github/lutsik/MeDeCom/src/R/utilities.R) to generate the cell-type proportions for pseudo-bulk samples. We followed the default parameter setup with the exception of 10 for proportion.var.factor and 1 million for number of genomic features. The numerical proportions of cell types in each pseudo-bulk sample is shown in **Supplementary Table 2**.

### 4.2 Differentially methylated regions

As gold-standard to be compared with informative region selection results showing cell-type-specific features, we extracted differentially methylated regions (DMRs). DMRs refer to genomic regions whose methylation states are different across given multiple samples. It is also known to be associated with the cell development and the cell reprogramming stages [59].

For mouse neuronal cell types, we compared each of 11 pure cell-type bulks (mL4, mL6-1, mL6-2, mDL-2, mL5-1, mL5-2, mL2-3, mSst-1, mNdnf-2, mPv and mVip) with all other bulks and identified DMRs respectively. For tumor analysis, we called DMRs between normal B-cell non-cancer and diffuse large B-cell lymphoma. All DMRs were called using DSS package 2.34.0 [60] with the following parameters: 0.2 for delta, 0.05 for threshold of p-values, 4 for minimum number of CpG sites and default values for minimum length and the distance to merge, which are 50 bps each.

### 4.3 CpG selection scheme for array-based deconvolution methods

Array-shaped methylome data comes in the matrix form, CpG sites by samples. To apply array-based deconvolution methods, Houseman and MeDeCom, as a comparison group, we transformed our sequencing data to array shape using *methrix* [61]. When converting the sequencing data, we specified a set of CpG sites to comprise the array shape. During data conversion for MeDeCom, we chose top 20000 CpGs with the highest beta-value variance. CpGs overlapping ctDMRs were taken to generate array-shaped data for Houseman’s method to adhere to the reference-based scenario.

### 4.4 Parameter values and procedures for each deconvolution method

We describe the detailed algorithm and parameter setting of each deconvolution method in this section. The pipelines are also available at https://github.com/CompEpigen/SeqDeconv_Pipeline.git.

#### 4.4.1 ClubCpG

ClubCpG [44] clusters reads fully covering informative regions satisfying given informative region selection conditions, using density-based spatial clustering of applications with noise (DBSCAN). For our experiments, we applied informative region selection conditions as given in **Table 1** which suited our dataset than the default values suggested by authors. Even though ClubCpG package itself does not have cell-type composition estimation function, we followed the cell-type deconvolution strategy proposed in [44]. Firstly, we created ClubCpG clustering results from another 100 pseudo-bulk samples as training data and extracted 20 principal components (PCs) using PCA algorithm from the result. Then, a multivariate linear regression model for cell-type proportion was fitted on the extracted 20 PCs. Finally, the cell-type composition was estimated using the trained multivariate linear regression model.

#### 4.4.2 PRISM

PRISM [42] infers the composition of epigenetically distinct subpopulations in tumor bulk samples based on methylation patterns. It mainly improves the accuracy by correcting erroneous methylation pattern using Hidden Markov Model. After the correction, the method remains only loci comprised of fully methylated and unmethylated patters, then cell-type proportions are estimated on the loci using EM algorithm. We applied PRISM with default setup as given in **Table 1** and yielded cell-type proportions in respective samples by calculating the ratio of inferred subclones.

#### 4.4.3 MethylPurify

MethylPurify [43] adopts EM algorithm to estimate tumor purity in bisulfite sequencing data. The EM algorithm in MethylPurify not only estimates methylation level of subpopulations, but also decides which subpopulation each read would be assigned to over iterations. We used the same parameter values in both mouse neuronal and tumor pseudo-bulks, 10 for read coverage and 50 for sampling time. However, bin length parameter was set to 300 bp for mouse neuronal pseudo-bulks and to 200 bp for tumor pseudo-bulks. Although the original code is designed to estimate cell-type compositions only within CpG islands, we altered it to conduct the estimation over all informative regions including non-CpG islands. This is because that the method initially failed in selecting enough number of informative regions for statistically significant cell-type composition estimation. After removing the CpG island filtering, we could obtain enough number of selected informative regions for cell-type deconvolution.

#### 4.4.4 csmFinder + coMethy

Yin et al. developed two distinct computational tools called csmFinder and coMethy [41]. csmFinder determines genomic regions showing cell-type specific signals denoted as putative cell-type-specific methylated (pCSM) loci. coMethy can decompose methylation level matrix of pCSM loci by samples, using NMF-approach similar to MeDeCom. For the specific input file format of csmFinder, we extracted methylation call results from our pseudo-bulk samples via *bismark_methylation_extractor* with *comprehensive, gzip* and *cytosine_report* options. CpG site coordinates of *hg19* and *mm10* genomes were used for corresponding samples. Running csmFinder, we followed the default parameters, minimum methylation difference of hypo- and hyper-methylation patterns 0.3 and p-value of the difference 0.05. Since coMethy works on a matrix which consists methylation patterns at same CpG sites from multiple samples, we collected pCSM loci detected from all samples in each experiment and filtered out only loci involving missing methylation values in any other sample than the sample that the locus was detected.

#### 4.4.5 BED

BED algorithm [40] is based on a Bayesian model and recognizes the distribution of epialleles which indicates all possible methylation patterns at CpG sites in a specific genomic range. For the preprocessing step to deal with contiguous and missing methylation patterns, we used pipelines elucidated in *bed-beta* Github page (https://github.com/james-e-barrett/bed-beta). This pipeline includes all epiallele estimation processes. Barret et al. demonstrated that the proportion of reads which are not attributed to normal tissue at each loci *i* can be calculated as follow:

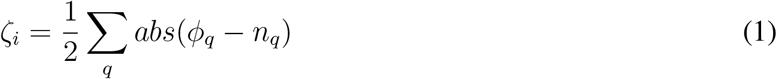

This equation is calculated with respect to all epialleles *q. ϕ* and *n* mean the distribution of epialleles at the locus and normal tissue samples each. We considered the rightmost maximum value in *ζ*_*i*_ distribution graph to be the estimated tumor purity as this is where the local epiallele distributions and normal tissue epiallele distributions showed the biggest discrepancy.

#### 4.4.6 DXM

DXM [46] is a computational method to infer not only the number of subpopulation within bulk methylomes but also the methylation profile of estimated subpopulations based on L1-norm minimization and Hidden Markov Model (HMM). It firstly investigates the distribution of methylation beta value over the given regions, and calculates L1-norm between the investigated distribution and 10,000 randomly generated distributions. The distribution with the lowest L1-norm is considered as an estimated cell-type distribution. Not likely other sequencing-based methods, DXM requires users to directly provide pre-selected specific genomic regions such as DMRs or CpG island regions. Therefore, keeping the reference-free manner and following the guidance of the authors, we gave methylation level within promoter and CpG Island regions as input of DXM after filtering the regions with minimum read coverage 4. CpG Island regions were downloaded from UCSC genome annotation database for mm10 and hg19 genomes (https://hgdownload.cse.ucsc.edu/goldenpath/mm10/database/ and https://hgdownload.cse.ucsc.edu/goldenpath/hg19/database/). For promoter regions, we also used UCSC annotation database exposed as TxDb objects [62, 63]. HMM is used mainly for the methylation profile inference, thus we do not include the details here.

#### 4.4.7 MeDeCom

MeDeCom [20] estimates cell-type proportions in array-shaped DNA methylation data based on nonnegative matrix factorization (NMF). When converting our sequencing-based data to array shape, we used *methrix* which is capable of loading *bedGraph* file format including methylation call and creates a CpG sites by samples matrix of methylation levels [61]. A bedGraph file for each pseudo-bulk sample was created through a computational tool called *MethylDackel* (https://github.com/dpryan79/MethylDackel). For reference genome to read bedGraph files in, *mm10* and *hg19* genomes were used for mouse neuronal and B-cell tumor pseudo-bulks each. As an input matrix of MeDeCom, we selected 20,000 CpG sites with the largest methylation level variance over covering reads. We ran MeDeCom with a regularization parameter *λ* from 10^−5^ to 10^−2^, 10 cross-validation folds, maximum iteration number 500 and random initialization number 30.

#### 4.4.8 Houseman’s method

Houseman’s method [15] (or Houseman short) was proposed to infer cell-type distribution from DNA methylation array-shaped data based on regression calibration. We used array-shaped data converted from our sequencing pseudo-bulk samples processed in the same way as for MeDeCom above. For selecting informative CpG sites, we mimicked the marker-CpG selection step of the original algorithm and chose CpG sites overlapping with ctDMRs which can provide cell-type specific signals. We increased the number of CpG sites to 1000 in the original code considering the larger number of total CpG sites in our pseudo-bulk dataset compared to microarray data they used. For the rest, we followed up the pipeline given in the supplementary material of the publication [15].

### 4.5 Performance measurement

#### 4.5.1 Genomic correlation

We used genomic correlation to calculate overall genomic base-wise proximity between informative region selection results and ctDMRs. This statistic was suggested by Favorov et al. for determinig the distribution of distances between two sets of genomic regions [49].

*Relative distance* of a selected informative region *q*_*i*_ with respect to given ctDMRs is defined as below:

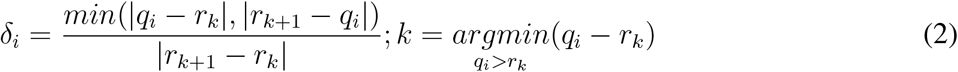

*r*_*k*_ and *r*_*k*+1_ are two nearest ctDMRs from given informative region *q*_*i*_. It is calculated by dividing the distance between selected informative region *q*_*i*_ and the closest DMR by the distance between two nearest ctDMRs.

If a set of selected informative regions and ctDMRs are independent, *δ*_*i*_ will have a uniform distribution. Hence, they test if the distribution of calculated relative distances makes a uniform distribution.

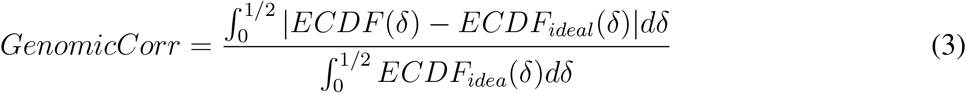

*ECDF* refers to empirical distirbution cumulative function and *ECDF* (*δ*) creates the distribution of observed *δ*_*i*_. *ECDF*_*ideal*_ is the uniform distribution. So, Equation 3 yields the ratio of difference between area under *ECDF* (*δ*) and under *ECDF*_*ideal*_. When given selected informative regions are independent of ctDMRs, *ECDF*_(_*δ*) will create a uniform distribution and *GenomicCorr* will reach 0. Conversely, if selected regions are identical to ctDMRs, *GenomicCorr* is 1.

#### 4.5.2 Absolute error between estimate and ground-truth

In cell-type composition estimation analysis, we assessed the performance by calculating absolute error between estimated proportion and ground-truth value of each bulk sample. For coMethy and MeDeCom, we had to match cell types to estimated components by choosing the one with the lowest mean absolute error out of all possible combinations of estimate and ground-truth pair.

#### 4.5.3 Entropy of cell-type distribution

In information theory, entropy is a concept to describe the level of uncertainty or information in a set. We used this measurement to determine how equally cell types are distributed in the cell mixture, because cell-type composition estimation can be more intractable with extremely low or high proportion of subpopulations. Applying entropy to our cell mixture, entropy of cell-type proportion in the cell mixture *C* comprised of *n* cell types, *c*_1_, *c*_2_, …, *c*_*n*_ is defined as:

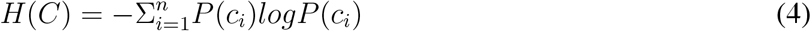

*P* (*c*_*i*_) refers to the cell-type proportion of cell type *c*_*i*_ here. Thus, the entropy value is higher when all cell-type proportions are more uniformly distributed.

For example, let us assume we have two cell mixtures *A* and *B* comprised of bi-components with different proportions. The dominant cell type takes apart of 90% in cell mixture *A* and 60% in cell mixture *B*. According to Equation 4, the entropy of cell mixture *A* is smaller than cell mixture *B*.

## Data availability

The single-cell mouse neuronal methylomes and tumour WGBS data are publicly available on NCBI GEO data repository (mouse neuronal single-cell: www.ncbi.nlm.nih.gov/geo/query/acc.cgi?acc=GSE97179, B-cell lymphoma and non-cancer WGBS: www.ncbi.nlm.nih.gov/geo/query/acc.cgi?acc=GSE137880). For the detailed data pre-processing, please see **Supplementary Table 1**.

## Code availability

All conducted analyses are reproducible following steps described in markdown files in our github repository (https://github.com/CompEpigen/SeqDeconv_Pipeline). Detailed piplines for each method are explained in the file named *METHOD_deconvolution_analysis*.*md*. We also uploaded the bug-fixed version of some methods through *git fork* function. The details of fixed lines in the code are clarified in *Commits* in each github repository. The list of methods uploaded as a new bug-fixed version is as below:

- **BED:** https://github.com/CompEpigen/bed-beta
- **DXM:** https://github.com/CompEpigen/dxm
- **MethylPurify:** https://github.com/CompEpigen/MethylPurify
- **PRISM:** https://github.com/CompEpigen/prism

For the methods we did not need to fix any bug, the code or the package is available here:

- **csmFinder:** https://github.com/Gavin-Yinld/csmFinder
- **coMethy:** https://github.com/Gavin-Yinld/coMethy
- **ClubCpG:** https://clubcpg.readthedocs.io/en/latest/
- **MeDeCom:** https://github.com/lutsik/MeDeCom
- **Houseman:** Supplementary file 2 on https://doi.org/10.1186/1471-2105-13-86

## Acknowledgement

We acknowledge guidance on their deconvolution method pipelines from James Barrett (UCL), Hehuang Xie (Virginia Tech), Robert A. Waterland (Baylor College of Medicine) and Anthony Scott (Mercury Data Science).

## Author contributions

Y.J mainly conducted all analyses and wrote the manuscript together with P.L. P.L also presented the idea, designed experiments and analyses together with Y.J. M.G contributed to the analysis result presentation. K.B and R.T contributed to data preprocessing pipelines. L.S and D.T contributed to the analyses conducted with DXM method. All authors reviewed the manuscript and provided critical feedback.

## Competing interests

The authors do not have competing interests.

## Notes

### Competing Interest Statement

The authors have declared no competing interest.

## References

[1] Omer Schwartzman and Amos Tanay. Single-cell epigenomics: techniques and emerging applications. Nature Reviews Genetics, 16(12):716–726, 2015.

[2] Peter Horak, Christoph Heining, Simon Kreutzfeldt, Barbara Hutter, Andreas Mock, Jennifer Hullein, Martina Frohlich, Sebastian Uhrig, Arne Jahn, Andreas Rump, et al. Comprehensive genomic and transcriptomic analysis for guiding therapeutic decisions in patients with rare cancers. Cancer Discovery, 2021.

[3] Katherine J Dick, Christopher P Nelson, Loukia Tsaprouni, Johanna K Sandling, Dylan Aïssi, Simone Wahl, Eshwar Meduri, Pierre-Emmanuel Morange, France Gagnon, Harald Grallert, et al. Dna methylation and body-mass index: a genome-wide analysis. The Lancet, 383(9933):1990–1998, 2014.

[4] Lucia L Lam, Eldon Emberly, Hunter B Fraser, Sarah M Neumann, Edith Chen, Gregory E Miller, and Michael S Kobor. Factors underlying variable dna methylation in a human community cohort. Proceedings of the National Academy of Sciences, 109(Supplement 2):17253–17260, 2012.

[5] ME Prince, R Sivanandan, A Kaczorowski, GT Wolf, MJ Kaplan, P Dalerba, IL Weissman, MF Clarke, and LE Ailles. Identification of a subpopulation of cells with cancer stem cell properties in head and neck squamous cell carcinoma. Proceedings of the National Academy of Sciences, 104(3):973–978, 2007.

[6] Yanhua Wen, Yanjun Wei, Shumei Zhang, Song Li, Hongbo Liu, Fang Wang, Yue Zhao, Dongwei Zhang, and Yan Zhang. Cell subpopulation deconvolution reveals breast cancer heterogeneity based on dna methylation signature. Briefings in bioinformatics, 18(3):426–440, 2017.

[7] Tony Hui, Qi Cao, Joanna Wegrzyn-Woltosz, Kieran O’Neill, Colin A Hammond, David JHF Knapp, Emma Laks, Michelle Moksa, Samuel Aparicio, Connie J Eaves, et al. High-resolution single-cell dna methylation measurements reveal epigenetically distinct hematopoietic stem cell subpopulations. Stem cell reports, 11(2):578–592, 2018.

[8] David Capper, David TW Jones, Martin Sill, Volker Hovestadt, Daniel Schrimpf, Dominik Sturm, Christian Koelsche, Felix Sahm, Lukas Chavez, David E Reuss, et al. Dna methylation-based classification of central nervous system tumours. Nature, 555(7697):469–474, 2018.

[9] Christian Koelsche, Daniel Schrimpf, Damian Stichel, Martin Sill, Felix Sahm, David E Reuss, Mirjam Blattner, Barbara Worst, Christoph E Heilig, Katja Beck, et al. Sarcoma classification by dna methylation profiling. Nature communications, 12(1):1–10, 2021.

[10] Alexey Kozlenkov, Minghui Wang, Panos Roussos, Sergei Rudchenko, Mihaela Barbu, Marina Bibikova, Brandy Klotzle, Andrew J Dwork, Bin Zhang, Yasmin L Hurd, et al. Substantial dna methylation differences between two major neuronal subtypes in human brain. Nucleic acids research, 44(6):2593–2612, 2016.

[11] Marco P Boks, Eske M Derks, Daniel J Weisenberger, Erik Strengman, Esther Janson, Iris E Sommer, Rene S Kahn, and Roel A Ophoff. The relationship of dna methylation with age, gender and genotype in twins and healthy controls. PloS one, 4(8):e6767, 2009.

[12] Fang Fang Zhang, Roberto Cardarelli, Joan Carroll, Kimberly G Fulda, Manleen Kaur, Karina Gonzalez, Jamboor K Vishwanatha, Regina M Santella, and Alfredo Morabia. Significant differences in global genomic dna methylation by gender and race/ethnicity in peripheral blood. Epigenetics, 6(5):623–629, 2011.

[13] Marina Bibikova, Bret Barnes, Chan Tsan, Vincent Ho, Brandy Klotzle, Jennie M Le, David Delano, Lu Zhang, Gary P Schroth, Kevin L Gunderson, et al. High density dna methylation array with single cpg site resolution. Genomics, 98(4):288–295, 2011.

[14] Ruth Pidsley, Elena Zotenko, Timothy J Peters, Mitchell G Lawrence, Gail P Risbridger, Peter Molloy, Susan Van Djik, Beverly Muhlhausler, Clare Stirzaker, and Susan J Clark. Critical evaluation of the illumina methylationepic beadchip microarray for whole-genome dna methylation profiling. Genome biology, 17(1):1–17, 2016.

[15] Eugene Andres Houseman, William P Accomando, Devin C Koestler, Brock C Christensen, Carmen J Marsit, Heather H Nelson, John K Wiencke, and Karl T Kelsey. Dna methylation arrays as surrogate measures of cell mixture distribution. BMC bioinformatics, 13(1):86, 2012.

[16] Ankur Chakravarthy, Andrew Furness, Kroopa Joshi, Ehsan Ghorani, Kirsty Ford, Matthew J Ward, Emma V King, Matt Lechner, Teresa Marafioti, Sergio A Quezada, et al. Pan-cancer deconvolution of tumour composition using dna methylation. Nature communications, 9(1):1–13, 2018.

[17] Andrew E Teschendorff, Charles E Breeze, Shijie C Zheng, and Stephan Beck. A comparison of reference-based algorithms for correcting cell-type heterogeneity in epigenome-wide association studies. BMC bioinformatics, 18(1):1–14, 2017.

[18] Hanyu Zhang, Ruoyi Cai, James Dai, and Wei Sun. Emeth: An em algorithm for cell type decomposition based on dna methylation data. Scientific reports, 11(1):1–12, 2021.

[19] Joshua J Levy, Alexander J Titus, Curtis L Petersen, Youdinghuan Chen, Lucas A Salas, and Brock C Christensen. Methylnet: an automated and modular deep learning approach for dna methylation analysis. BMC bioinformatics, 21(1):1–15, 2020.

[20] Pavlo Lutsik, Martin Slawski, Gilles Gasparoni, Nikita Vedeneev, Matthias Hein, and Jörn Walter. Medecom: discovery and quantification of latent components of heterogeneous methylomes. Genome biology, 18(1):1–20, 2017.

[21] E Andres Houseman, Molly L Kile, David C Christiani, Tan A Ince, Karl T Kelsey, and Carmen J Marsit. Reference-free deconvolution of dna methylation data and mediation by cell composition effects. BMC bioinformatics, 17(1):1–15, 2016.

[22] Vitor Onuchic, Ryan J Hartmaier, David N Boone, Michael L Samuels, Ronak Y Patel, Wendy M White, Vesna D Garovic, Steffi Oesterreich, Matt E Roth, Adrian V Lee, et al. Epigenomic deconvolution of breast tumors reveals metabolic coupling between constituent cell types. Cell reports, 17(8):2075–2086, 2016.

[23] Elior Rahmani, Regev Schweiger, Liat Shenhav, Theodora Wingert, Ira Hofer, Eilon Gabel, Eleazar Eskin, and Eran Halperin. Bayescce: a bayesian framework for estimating cell-type composition from dna methylation without the need for methylation reference. Genome biology, 19(1):1–18, 2018.

[24] Elior Rahmani, Regev Schweiger, Brooke Rhead, Lindsey A Criswell, Lisa F Barcellos, Eleazar Eskin, Saharon Rosset, Sriram Sankararaman, and Eran Halperin. Cell-type-specific resolution epigenetics without the need for cell sorting or single-cell biology. Nature communications, 10(1):1–11, 2019.

[25] Michael Scherer, Florian Schmidt, Olga Lazareva, Jörn Walter, Jan Baumbach, Marcel H Schulz, and Markus List. Machine learning for deciphering cell heterogeneity and gene regulation. Nature Computational Science, 1(3):183–191, 2021.

[26] Clémentine Decamps, Florian Privé, Raphael Bacher, Daniel Jost, Arthur Waguet, Eugene Andres Houseman, Eugene Lurie, Pavlo Lutsik, Aleksandar Milosavljevic, Michael Scherer, et al. Guidelines for cell-type heterogeneity quantification based on a comparative analysis of reference-free dna methylation deconvolution software. BMC bioinformatics, 21(1):1–15, 2020.

[27] Benjamin Goeppert, Reka Toth, Stephan Singer, Thomas Albrecht, Daniel B Lipka, Pavlo Lutsik, David Brocks, Marion Baehr, Oliver Muecke, Yassen Assenov, et al. Integrative analysis defines distinct prognostic subgroups of intrahepatic cholangiocarcinoma. Hepatology, 69(5):2091–2106, 2019.

[28] Michael Scherer, Petr V Nazarov, Reka Toth, Shashwat Sahay, Tony Kaoma, Valentin Maurer, Nikita Vedeneev, Christoph Plass, Thomas Lengauer, Jörn Walter, et al. Reference-free deconvolution, visualization and interpretation of complex dna methylation data using decomppipeline, medecom and factorviz. Nature Protocols, 15(10):3240–3263, 2020.

[29] Yuanyuan Chen, Reka Toth, Sara Chocarro, Dieter Weichenhan, Joschka Hey, Pavlo Lutsik, Stefan Sawall, Georgios T Stathopoulos, Christoph Plass, and Rocio Sotillo. Diverse routes of club cell evolution in lung adenocarcinoma. bioRxiv, 2021.

[30] Malte Simon, Sadaf S Mughal, Peter Horak, Sebastian Uhrig, Jonas Buchloh, Bogac Aybey, Albrecht Stenzinger, Hanno Glimm, Stefan Fröhling, Benedikt Brors, et al. Deconvolution of sarcoma methylomes reveals varying degrees of immune cell infiltrates with association to genomic aberrations. Journal of translational medicine, 19(1):1–17, 2021.

[31] Alexander Meissner, Andreas Gnirke, George W Bell, Bernard Ramsahoye, Eric S Lander, and Rudolf Jaenisch. Reduced representation bisulfite sequencing for comparative high-resolution dna methylation analysis. Nucleic acids research, 33(18):5868–5877, 2005.

[32] Ryan Lister, Mattia Pelizzola, Robert H Dowen, R David Hawkins, Gary Hon, Julian Tonti-Filippini, Joseph R Nery, Leonard Lee, Zhen Ye, Que-Minh Ngo, et al. Human dna methylomes at base resolution show widespread epigenomic differences. nature, 462(7271):315–322, 2009.

[33] Chang Shu, Xinyu Zhang, Bradley E Aouizerat, and Ke Xu. Comparison of methylation capture sequencing and infinium methylationepic array in peripheral blood mononuclear cells. Epigenetics & chromatin, 13(1):1–15, 2020.

[34] Wanding Zhou, Huy Q Dinh, Zachary Ramjan, Daniel J Weisenberger, Charles M Nicolet, Hui Shen, Peter W Laird, and Benjamin P Berman. Dna methylation loss in late-replicating domains is linked to mitotic cell division. Nature genetics, 50(4):591–602, 2018.

[35] Abdulrahman Salhab, Karl Nordström, Gilles Gasparoni, Kathrin Kattler, Peter Ebert, Fidel Ramirez, Laura Arrigoni, Fabian Müller, Julia K Polansky, Cristina Cadenas, et al. A comprehensive analysis of 195 dna methylomes reveals shared and cell-specific features of partially methylated domains. Genome biology, 19(1):1–13, 2018.

[36] Stephen J Clark, Sébastien A Smallwood, Heather J Lee, Felix Krueger, Wolf Reik, and Gavin Kelsey. Genome-wide base-resolution mapping of dna methylation in single cells using single-cell bisulfite sequencing (scbs-seq). Nature protocols, 12(3):534–547, 2017.

[37] Hongshan Guo, Ping Zhu, Xinglong Wu, Xianlong Li, Lu Wen, and Fuchou Tang. Single-cell methylome landscapes of mouse embryonic stem cells and early embryos analyzed using reduced representation bisulfite sequencing. Genome research, 23(12):2126–2135, 2013.

[38] Ricard Argelaguet, Stephen J Clark, Hisham Mohammed, L Carine Stapel, Christel Krueger, Chantriolnt-Andreas Kapourani, Ivan Imaz-Rosshandler, Tim Lohoff, Yunlong Xiang, Courtney W Hanna, et al. Multi-omics profiling of mouse gastrulation at single-cell resolution. Nature, 576(7787):487–491, 2019.

[39] Shuhui Bian, Yu Hou, Xin Zhou, Xianlong Li, Jun Yong, Yicheng Wang, Wendong Wang, Jia Yan, Boqiang Hu, Hongshan Guo, et al. Single-cell multiomics sequencing and analyses of human colorectal cancer. Science, 362(6418):1060–1063, 2018.

[40] James E Barrett, Andrew Feber, Javier Herrero, Miljana Tanic, Gareth A Wilson, Charles Swanton, and Stephan Beck. Quantification of tumour evolution and heterogeneity via bayesian epiallele detection. BMC bioinformatics, 18(1):1–10, 2017.

[41] Liduo Yin, Yanting Luo, Xiguang Xu, Shiyu Wen, Xiaowei Wu, Xuemei Lu, and Hehuang Xie. Virtual methylome dissection facilitated by single-cell analyses. Epigenetics & chromatin, 12(1):1–13, 2019.

[42] Dohoon Lee, Sangseon Lee, and Sun Kim. Prism: methylation pattern-based, reference-free inference of subclonal makeup. Bioinformatics, 35(14):i520–i529, 2019.

[43] Xiaoqi Zheng, Qian Zhao, Hua-Jun Wu, Wei Li, Haiyun Wang, Clifford A Meyer, Qian Alvin Qin, Han Xu, Chongzhi Zang, Peng Jiang, et al. Methylpurify: tumor purity deconvolution and differential methylation detection from single tumor dna methylomes. Genome biology, 15(7):1–13, 2014.

[44] C Anthony Scott, Jack D Duryea, Harry MacKay, Maria S Baker, Eleonora Laritsky, Chathura J Gunasekara, Cristian Coarfa, and Robert A Waterland. Identification of cell type-specific methylation signals in bulk whole genome bisulfite sequencing data. Genome biology, 21(1):1–23, 2020.

[45] Alexander J Titus, Rachel M Gallimore, Lucas A Salas, and Brock C Christensen. Cell-type deconvolution from dna methylation: a review of recent applications. Human molecular genetics, 26(R2):R216–R224, 2017.

[46] Jerry Fong, Jacob R Gardner, Jared M Andrews, Amanda F Cashen, Jacqueline E Payton, Kilian Q Weinberger, and John R Edwards. Determining subpopulation methylation profiles from bisulfite sequencing data of heterogeneous samples using dxm. Nucleic acids research, 49(16):e93–e93, 2021.

[47] Chongyuan Luo, Christopher L Keown, Laurie Kurihara, Jingtian Zhou, Yupeng He, Junhao Li, Rosa Castanon, Jacinta Lucero, Joseph R Nery, Justin P Sandoval, et al. Single-cell methylomes identify neuronal subtypes and regulatory elements in mammalian cortex. Science, 357(6351):600–604, 2017.

[48] Catherine Do, Emmanuel LP Dumont, Martha Salas, Angelica Castano, Huthayfa Mujahed, Leonel Maldonado, Arunjot Singh, Sonia C DaSilva-Arnold, Govind Bhagat, Soren Lehman, et al. Allelespecific dna methylation is increased in cancers and its dense mapping in normal plus neoplastic cells increases the yield of disease-associated regulatory snps. Genome biology, 21(1):1–39, 2020.

[49] Alexander Favorov, Loris Mularoni, Leslie M Cope, Yulia Medvedeva, Andrey A Mironov, Vsevolod J Makeev, and Sarah J Wheelan. Exploring massive, genome scale datasets with the genometricorr package. PLoS Comput Biol, 8(5):e1002529, 2012.

[50] Michael T McCabe, Johann C Brandes, and Paula M Vertino. Cancer dna methylation: molecular mechanisms and clinical implications. Clinical Cancer Research, 15(12):3927–3937, 2009.

[51] Yang Liu, Shuai He, Xi-Liang Wang, Wan Peng, Qiu-Yan Chen, Dong-Mei Chi, Jie-Rong Chen, Bo-Wei Han, Guo-Wang Lin, Yi-Qi Li, et al. Tumour heterogeneity and intercellular networks of nasopharyngeal carcinoma at single cell resolution. Nature communications, 12(1):1–18, 2021.

[52] Xiao Dong, Fan Wang, Chuan Liu, Jing Ling, Xuebing Jia, Feifei Shen, Ning Yang, Sibo Zhu, Lin Zhong, and Qi Li. Single-cell analysis reveals the intra-tumor heterogeneity and identifies mlxipl as a biomarker in the cellular trajectory of hepatocellular carcinoma. Cell death discovery, 7(1):1–13, 2021.

[53] Yan Zhou, Dong Yang, Qingcheng Yang, Xiaobin Lv, Wentao Huang, Zhenhua Zhou, Yaling Wang, Zhichang Zhang, Ting Yuan, Xiaomin Ding, et al. Single-cell rna landscape of intratumoral heterogeneity and immunosuppressive microenvironment in advanced osteosarcoma. Nature communications, 11(1):1–17, 2020.

[54] Roghayyeh Baghban, Leila Roshangar, Rana Jahanban-Esfahlan, Khaled Seidi, Abbas Ebrahimi-Kalan, Mehdi Jaymand, Saeed Kolahian, Tahereh Javaheri, and Peyman Zare. Tumor microenvironment complexity and therapeutic implications at a glance. Cell Communication and Signaling, 18(1):1–19, 2020.

[55] Felix Krueger and Simon R Andrews. Bismark: a flexible aligner and methylation caller for bisulfite-seq applications. bioinformatics, 27(11):1571–1572, 2011.

[56] Marcel Martin. Cutadapt removes adapter sequences from high-throughput sequencing reads. EMBnet. journal, 17(1):10–12, 2011.

[57] Heng Li, Bob Handsaker, Alec Wysoker, Tim Fennell, Jue Ruan, Nils Homer, Gabor Marth, Goncalo Abecasis, and Richard Durbin. The sequence alignment/map format and samtools. Bioinformatics, 25(16):2078–2079, 2009.

[58] Catherine Do, Charles F Lang, John Lin, Huferesh Darbary, Izabela Krupska, Aulona Gaba, Lynn Petukhova, Jean-Paul Vonsattel, Mary P Gallagher, Robin S Goland, et al. Mechanisms and disease associations of haplotype-dependent allele-specific dna methylation. The American Journal of Human Genetics, 98(5):934–955, 2016.

[59] Michel Neidhart. DNA methylation and complex human disease. Academic Press, 2015.

[60] Hao Wu, Chi Wang, and Zhijin Wu. A new shrinkage estimator for dispersion improves differential expression detection in rna-seq data. Biostatistics, 14(2):232–243, 2013.

[61] Anand Mayakonda, Maximilian Schönung, Joschka Hey, Rajbir Nath Batra, Clarissa Feuerstein-Akgoz, Kristin Köhler, Daniel B Lipka, Rocio Sotillo, Christoph Plass, Pavlo Lutsik, and Reka Toth. Methrix: an R/Bioconductor package for systematic aggregation and analysis of bisulfite sequencing data. Bioinformatics, 36(22-23):5524–5525, 12 2020.

[62] BC Team and BP Maintainer. Txdb. mmusculus. ucsc. mm10. knowngene: Annotation package for txdb object (s). r package version 3.10. 0. 2020.

[63] Marc Carlson and Bioconductor Package Maintainer. Txdb. hsapiens. ucsc. hg19. knowngene. 2015.

