## Supplementary for "Systematic evaluation of cell-type deconvolution pipelines for sequencing-based bulk DNA methylomes"

| Dataset | Step | Tool | Command line |
|---|---|---|---|
| **GSE97179** *Mouse neuronal scBS-seq* | Trimming 1 | Cutadapt 2.6 | cutadapt -f fastq -q 20 -m 62 -a AGATCGGAAGAGCACACGTCTGAAC -A AGATCGGAAGAGCGTCGTGTAGGGA |
| | Trimming 2 | Cutadapt 2.6 | cutadapt -f fastq -u 16 -u -16 -U 16 -U -16 -m 30 |
| | Alignment | Bismark 0.22.3 | bismark –bowtie2 –se (–pbat only for the first reads) |
| | Sorting | Samtools 1.9 | samtools sort |
| | Duplicate removal | picard MarkDuplicates 1.141 | picard MarkDuplicates REMOVE DUPLICATES=true |
| | Filtering | Samtools 1.9 | samtools view -b -q 30 |
| **GSE137880** *B cell tumor WGBS* | Trimming | Trim Galore 0.6.6 | trim galore -q 30 –length 40 –paired –retain unpair |
| | Alignment 1 | Bismark 0.22.3 | bismark –bowtie2 |
| | Alignment 2 | Bismark 0.22.3 | bismark –bowtie2 -non_directional |
| | Sorting | Samtools 1.9 | samtools sort |
| | Duplicate removal | picard MarkDuplicates 1.141 | picard MarkDuplicates REMOVE DUPLICATES=true |

**Table 1** Preprocessing steps for mouse neuronal scBS-seq and B cell WGBS data. Each step is clarified with a specific version of tool we used for our preprocessing. Trimming 2 step in mouse neuronal scBS-seq data precessing is required only for multiplexed samples according to Luo et. al. [1] and the alignment 2 step in B cell tumor WGBS preprocessing was done only for the unmatched reads after the paired-end alignment during the alignment 1.

| | 2 cell-type mouse neuron | | 5 cell-type mouse neuron | | | | | Tumor | |
|---|---|---|---|---|---|---|---|---|---|
| | mL6-2 | mPv | mDL-2 | mL2-3 | mL5-1 | mL6-2 | mPv | B cell non-cancer | B cell Lymphoma |
| **Bulk 1** | 0.731 | 0.269 | 0.350 | 0.148 | 0.242 | 0.091 | 0.169 | 0.151 | 0.849 |
| **Bulk 2** | 0.445 | 0.555 | 0.149 | 0.096 | 0.451 | 0.106 | 0.197 | 0.945 | 0.055 |
| **Bulk 3** | 0.810 | 0.190 | 0.444 | 0.166 | 0.078 | 0.293 | 0.020 | 0.152 | 0.848 |
| **Bulk 4** | 0.658 | 0.342 | 0.376 | 0.176 | 0.245 | 0.091 | 0.112 | 0.190 | 0.810 |
| **Bulk 5** | 0.338 | 0.662 | 0.049 | 0.459 | 0.062 | 0.172 | 0.258 | 0.680 | 0.320 |
| **Bulk 6** | 0.352 | 0.648 | 0.381 | 0.100 | 0.037 | 0.407 | 0.074 | 0.801 | 0.199 |
| **Bulk 7** | 0.617 | 0.383 | 0.176 | 0.035 | 0.242 | 0.317 | 0.230 | 0.790 | 0.210 |
| **Bulk 8** | 0.591 | 0.409 | 0.141 | 0.075 | 0.124 | 0.249 | 0.410 | 0.496 | 0.504 |
| **Bulk 9** | 0.558 | 0.442 | 0.160 | 0.199 | 0.166 | 0.151 | 0.324 | 0.624 | 0.376 |
| **Bulk 10** | 0.444 | 0.556 | 0.280 | 0.130 | 0.456 | 0.034 | 0.100 | 0.657 | 0.343 |
| **Bulk 11** | 0.330 | 0.669 | 0.259 | 0.155 | 0.264 | 0.290 | 0.032 | 0.552 | 0.448 |
| **Bulk 12** | 0.377 | 0.623 | 0.081 | 0.446 | 0.140 | 0.166 | 0.166 | 0.963 | 0.037 |
| **Bulk 13** | 0.662 | 0.338 | 0.248 | 0.142 | 0.141 | 0.118 | 0.351 | 0.955 | 0.045 |
| **Bulk 14** | 0.461 | 0.539 | 0.119 | 0.340 | 0.118 | 0.245 | 0.177 | 0.315 | 0.685 |
| **Bulk 15** | 0.835 | 0.165 | 0.150 | 0.062 | 0.278 | 0.295 | 0.215 | 0.983 | 0.017 |
| **Bulk 16** | 0.694 | 0.306 | 0.201 | 0.451 | 0.088 | 0.241 | 0.019 | 0.804 | 0.196 |
| **Bulk 17** | 0.550 | 0.450 | 0.266 | 0.210 | 0.210 | 0.169 | 0.146 | 0.170 | 0.830 |
| **Bulk 18** | 0.624 | 0.376 | 0.350 | 0.027 | 0.340 | 0.225 | 0.058 | 0.738 | 0.262 |
| **Bulk 19** | 0.671 | 0.329 | 0.271 | 0.159 | 0.334 | 0.198 | 0.039 | 0.673 | 0.327 |
| **Bulk 20** | 0.539 | 0.461 | 0.053 | 0.390 | 0.055 | 0.112 | 0.390 | 0.926 | 0.074 |

**Table 2** Cell-type composition of each sample in three pseudo-bulk datasets. We generated 20 pseudo-bulk samples for individual datasets which are grouped by thick vertical lines in the table. The compositions are drawn from Dirichlet distribution as explained in Method.

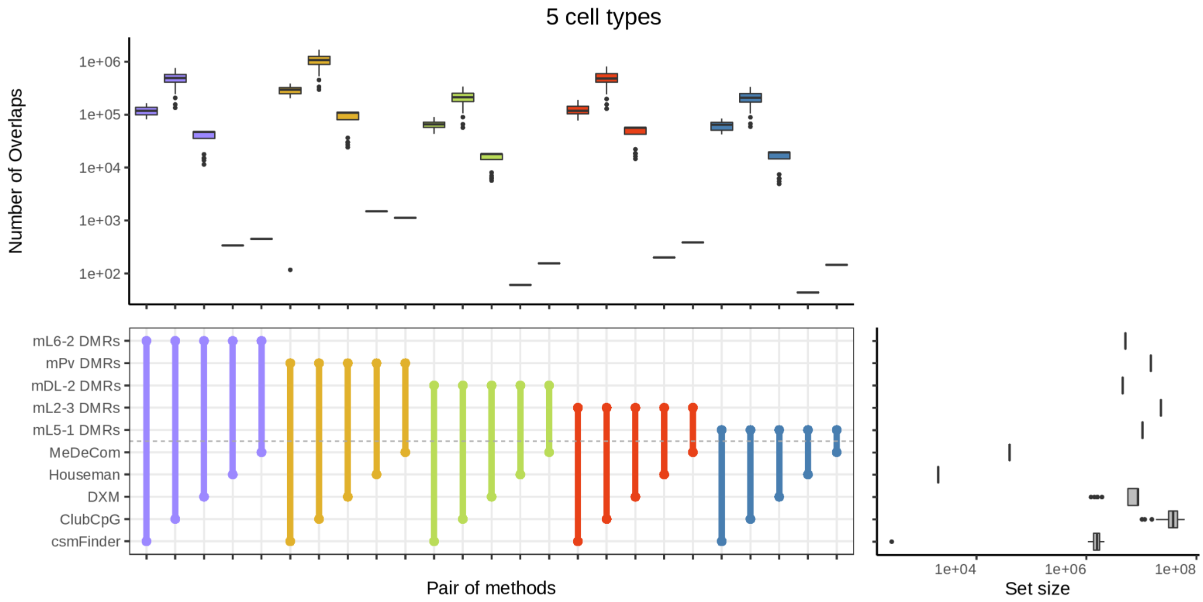


**Figure 1:** Overlaps between cell-type DMRs and selected informative regions for five cell-type mouse neuronal pseudo-bulks. THe colored box plots at the top shows the number of overlaps between a pair of methods connected at the middle, across all methods. The grey box plot at the right side means the number of informative regions detected by each method or the number of regions in DMRs.

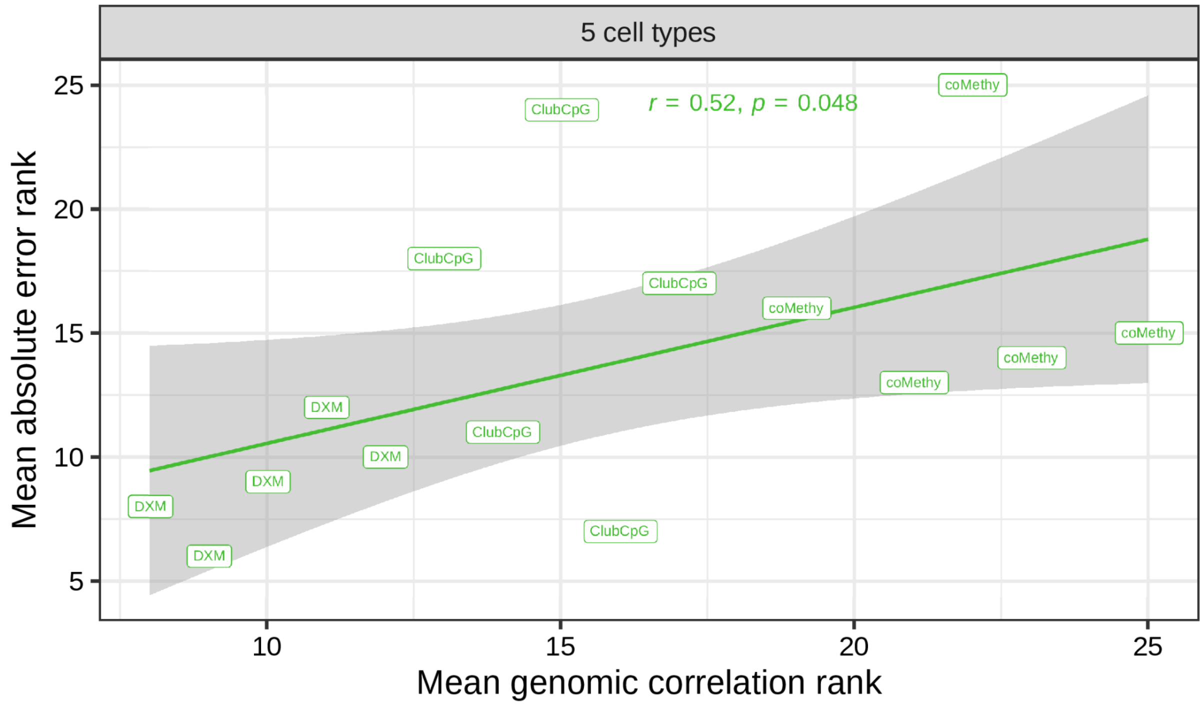


**Figure 2:** Mean absolute error vs mean genomic correlation between selected informative regions and DMRs in five cell-type mouse neuronal pseudo-bulks. The points indicate results of each cell type from each deconvolution method.

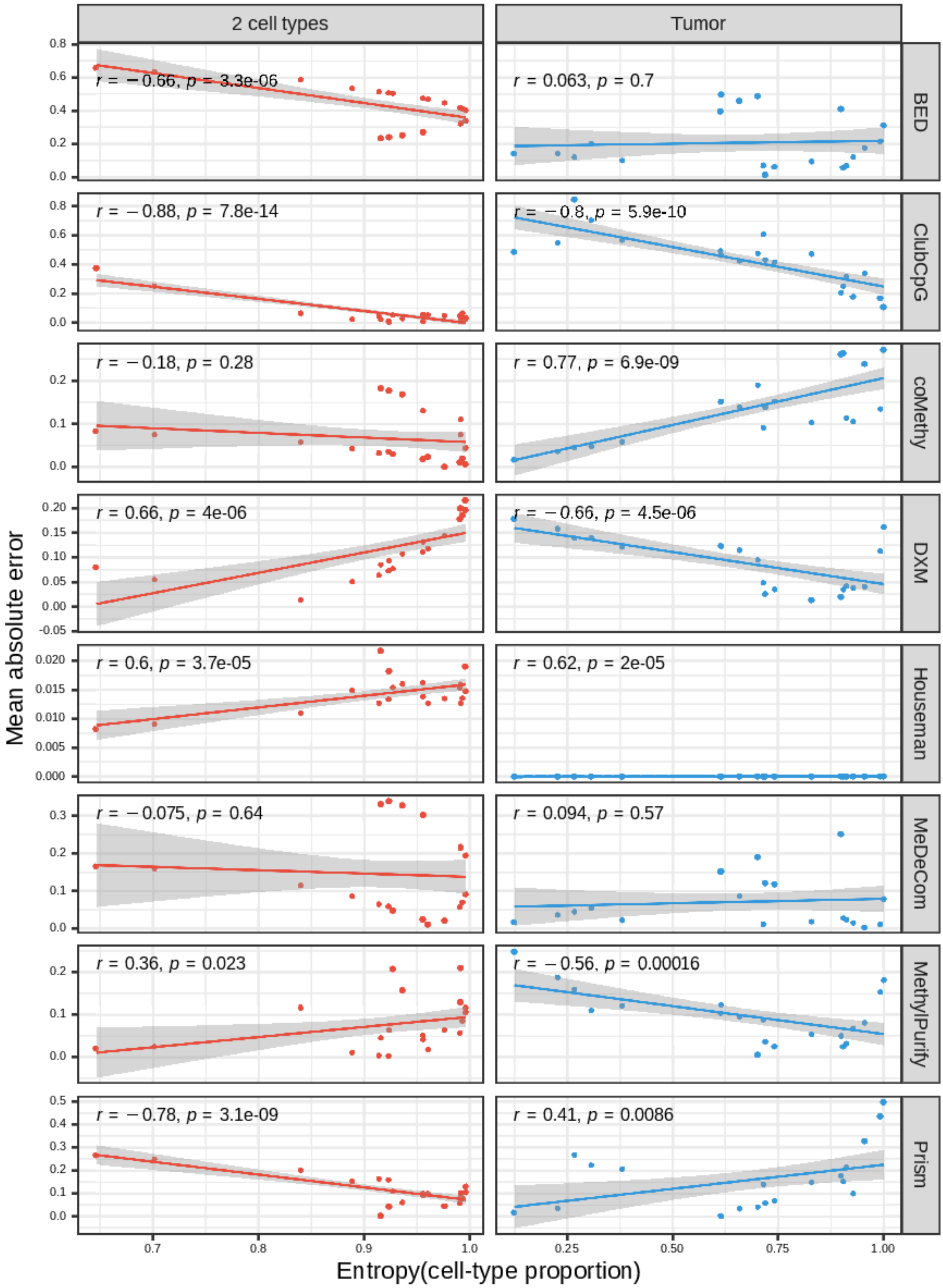


**Figure 3** Mean absolute error vs entropy of cell-type proportions in each bulk sample (2 cell-type mouse neuronal and tumor pseudo-bulks). Dots are fitted in a linear function (the line with grey background).

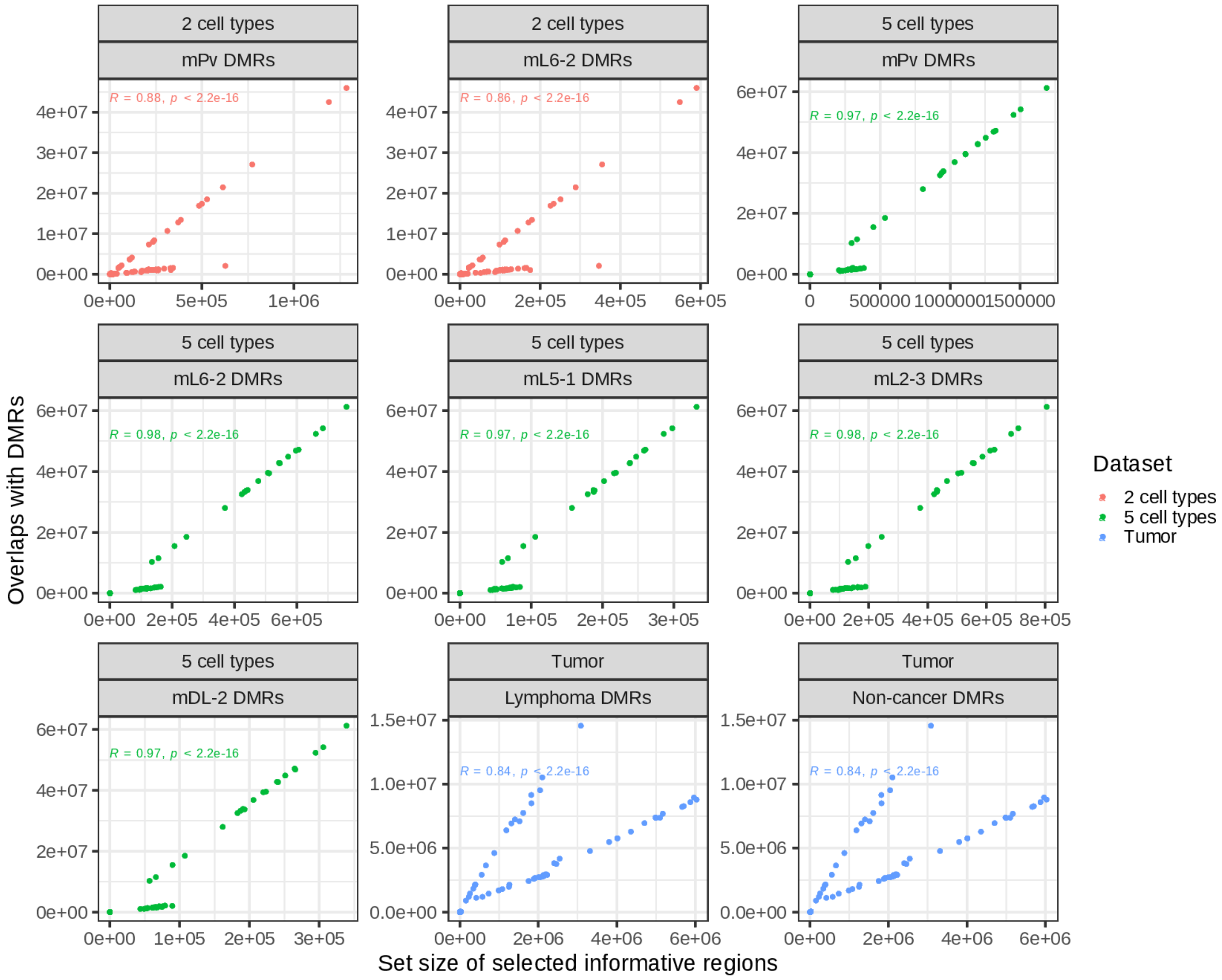


**Figure 4** Correlation between the number of selected informative regions and overlaps with DMRs. Each point presents a result by each method in each bulk. Different colors are applied to respective datasets.

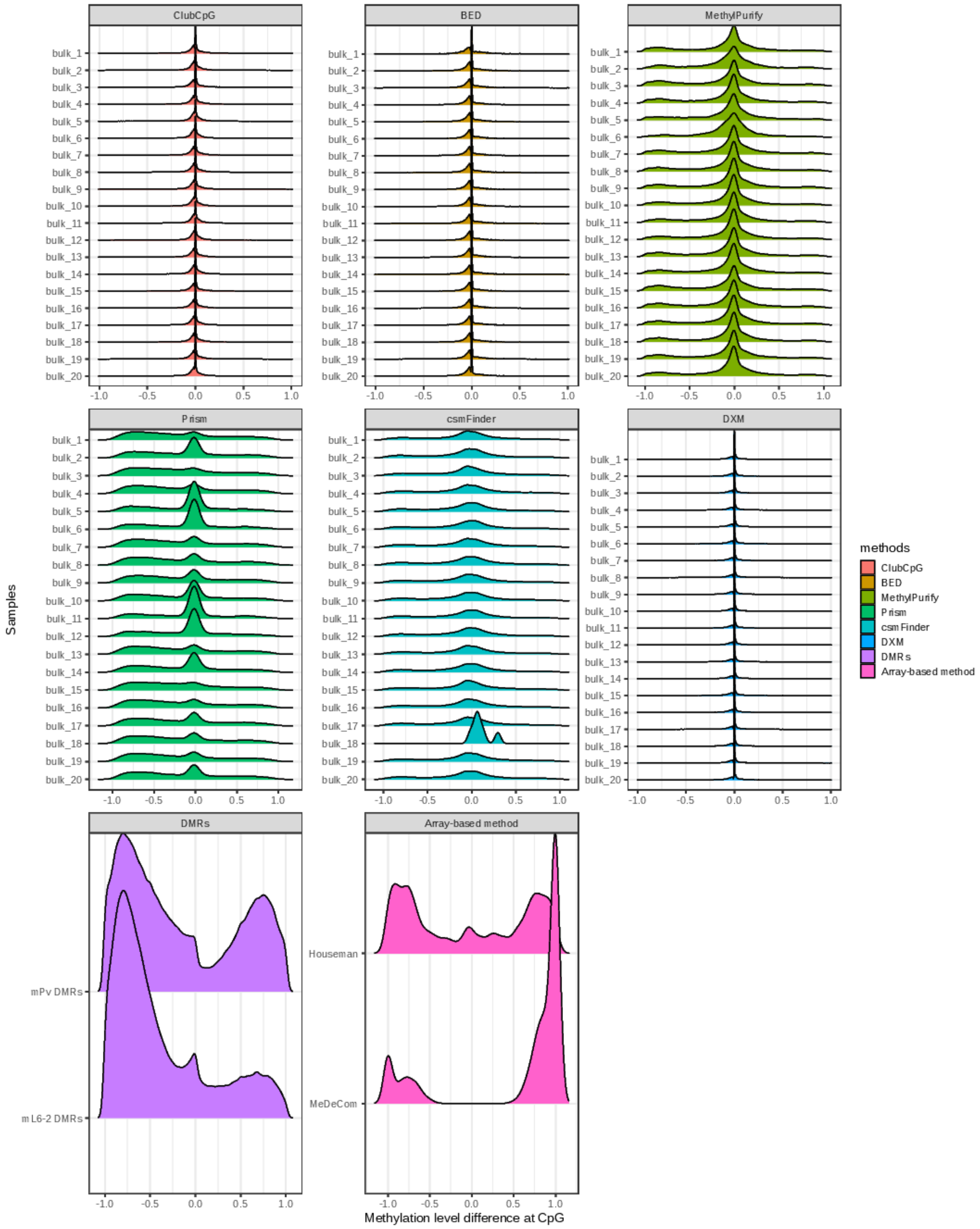


**Figure 5** Difference of methylation beta values between targeted cell types for 2 cell-type mouse neuronal pseudo-bulks (mL6-2 and mPv) within selected informative regions by each method. We calculated the difference of methylation beta values in CpGs overlapping with the selected informative regions and this figure presents the distribution of differences by samples and by methods. Since array-based methods use same informative regions for all samples, each method has one distribution.

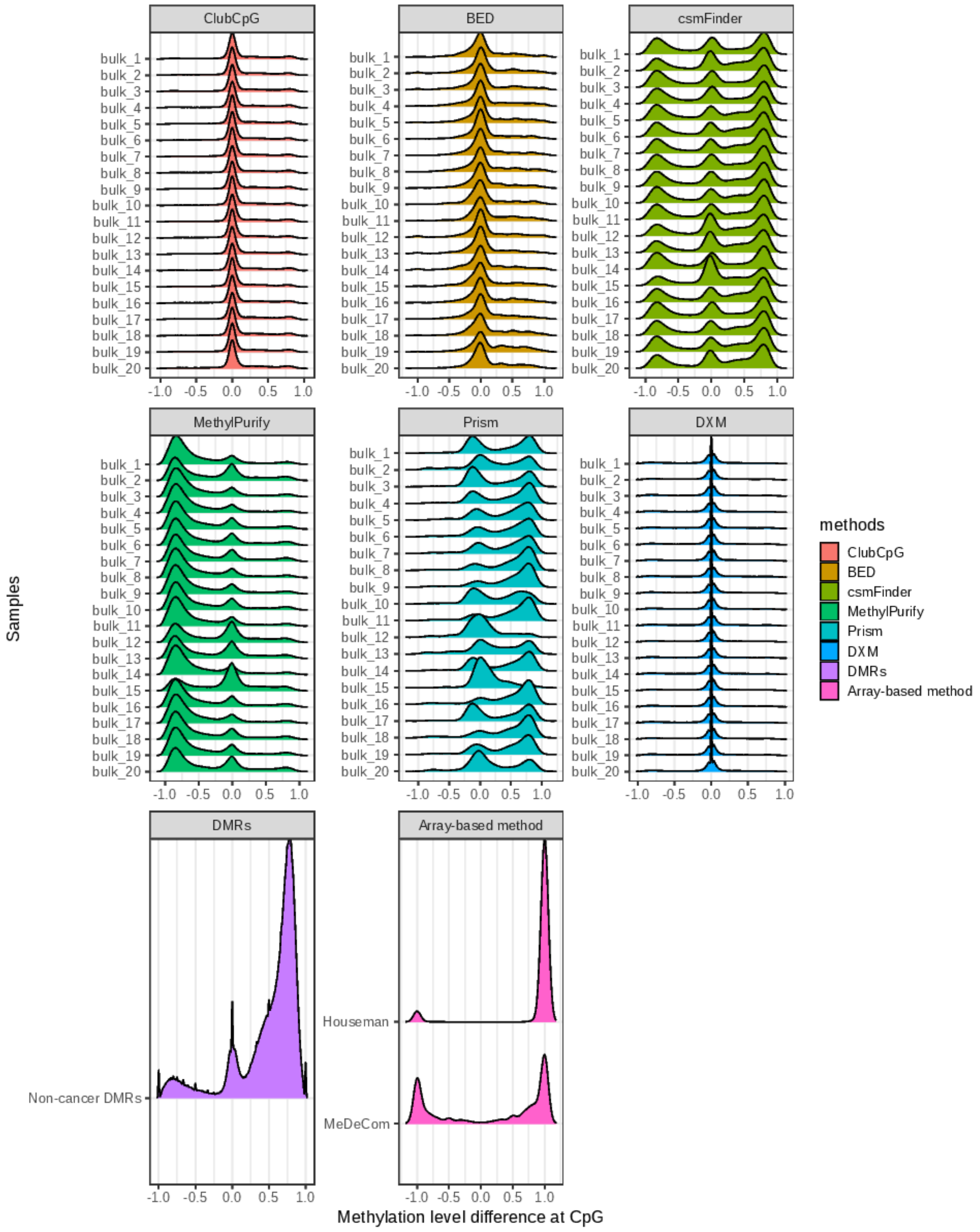


**Figure 6** Difference of methylation beta values between targeted cell types for tumor pseudo-bulks (B cell non-cancer and B cell lymphoma) within selected informative regions by each method. We calculated the difference of methylation beta values in CpGs overlapping with the selected informative regions and this figure presents the distribution of differences by samples and by methods. Since array-based methods use same informative regions for all samples, each method has one distribution. B cell non-cancer and lymphoma DMRs, yielded by comparing only those two cell types, are equal each other, thus we included only B cell non-cancer DMRs in the result.

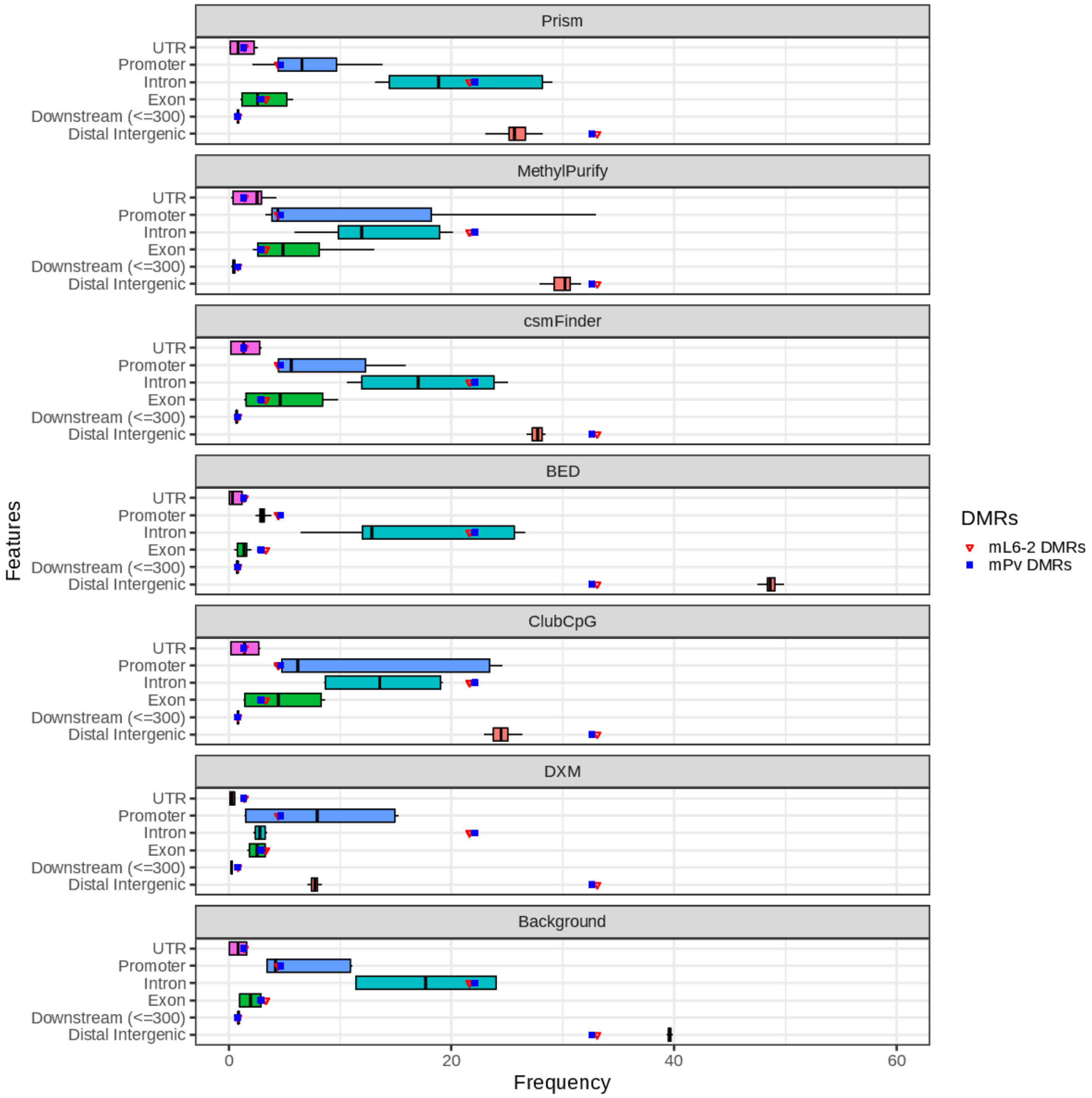


**Figure 7** Peak annotation analysis on selected informative regions by each method in 2 cell-type mouse neuronal pseudo-bulks. Analyses were done separately in respective bulks, and the frequency is presented in a box plot with median value (the middle line), the first and the third quantiles (the ends of box). Peak annotations within DMRs are presented as dots in each box plot. For the background, we conducted peak annotation over the entire CpG sites of each bulk.

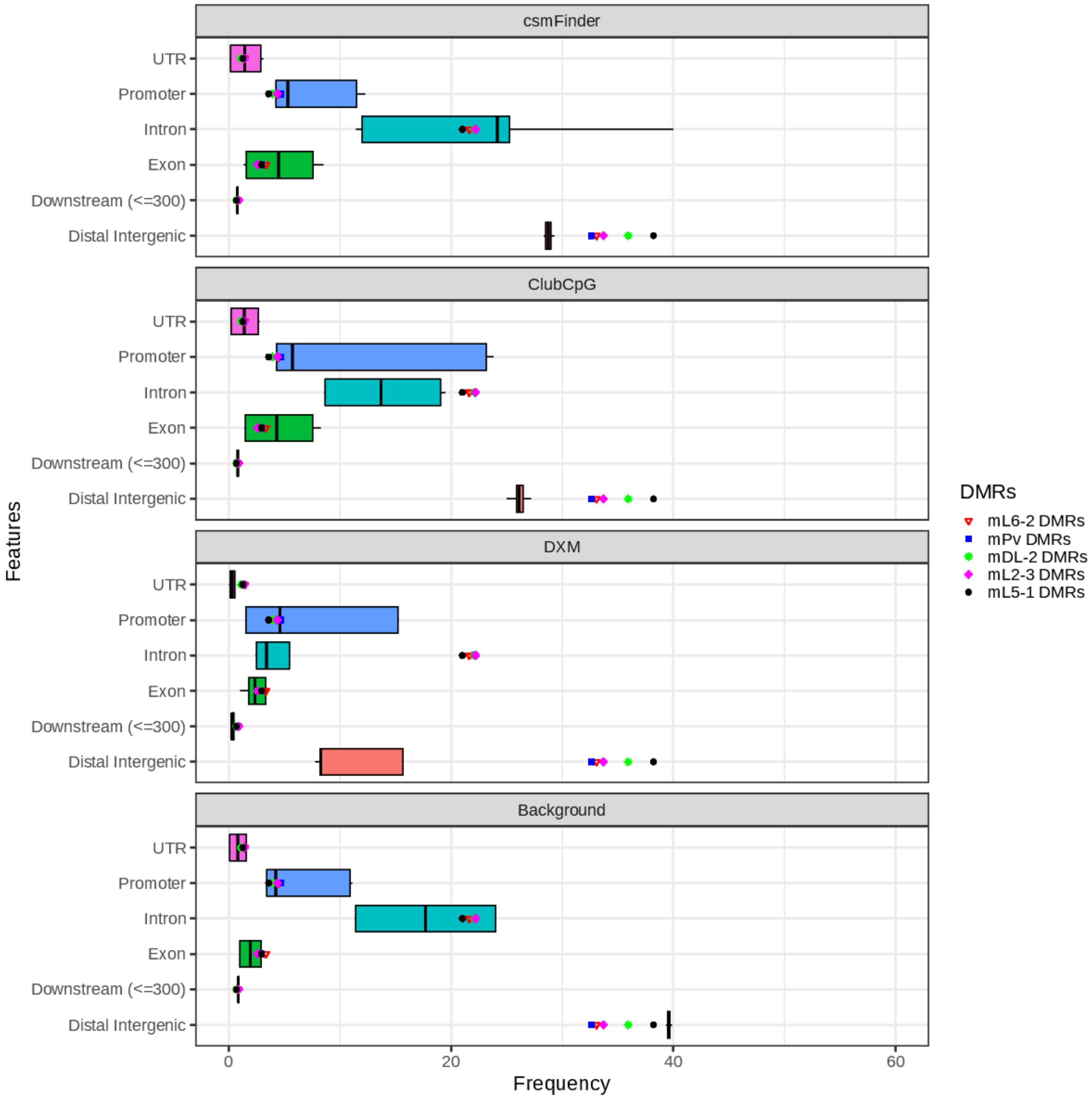


**Figure 8** Peak annotation analysis on selected informative regions by each method in 5 cell-type mouse neuronal pseudo-bulks. Analyses were done separately in respective bulks, and the frequency is presented in a box plot with median value (the middle line), the first and the third quantiles (the ends of box). Peak annotations within DMRs are presented as dots in each box plot. For the background, we conducted peak annotation over the entire CpG sites of each bulk.

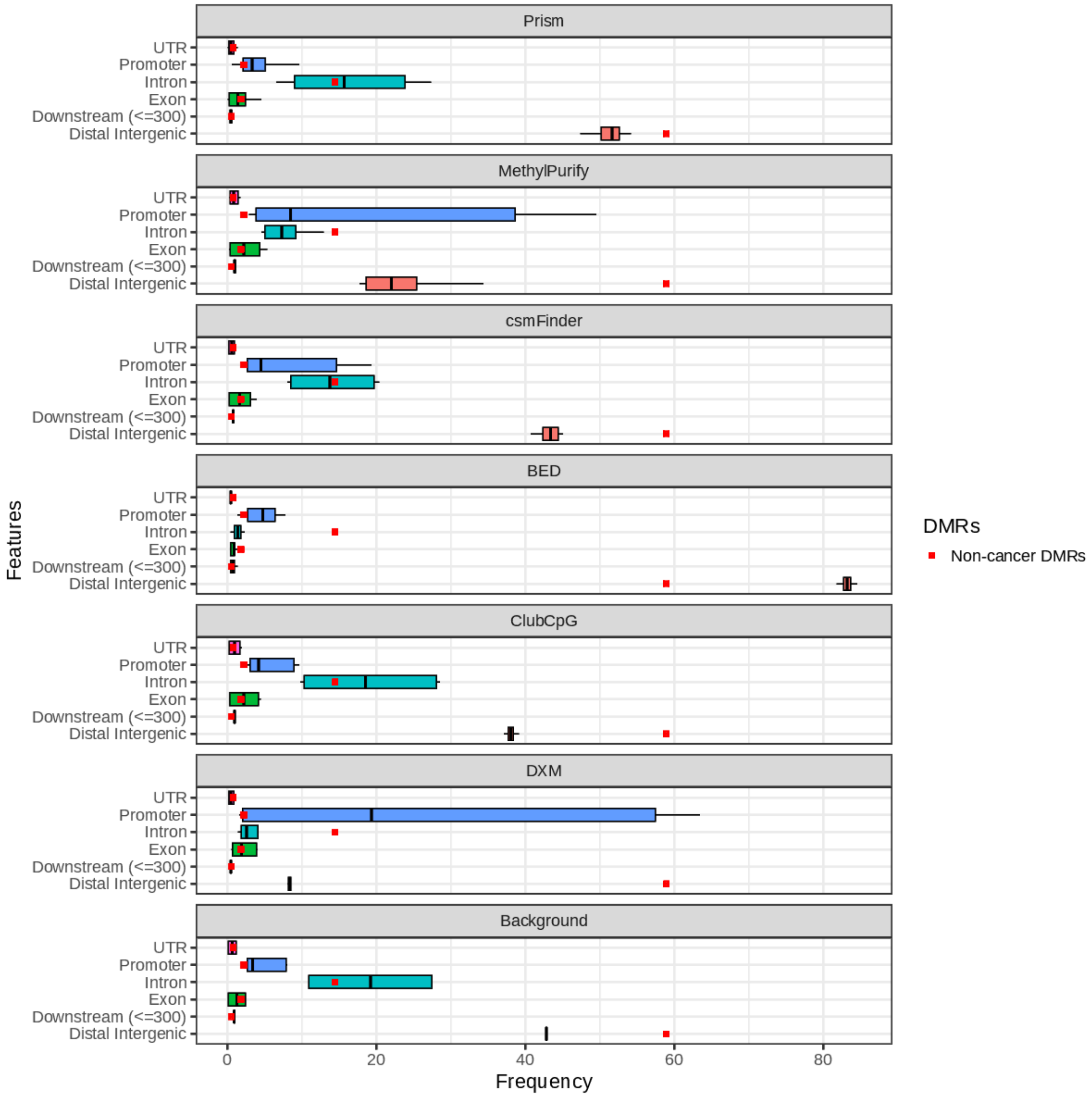


**Figure 9** Peak annotation analysis on selected informative regions by each method in tumor pseudo-bulks. Analyses were done separately in respective bulks, and the frequency is presented in a box plot with median value (the middle line), the first and the third quantiles (the ends of box). Peak annotations within DMRs are presented as dots in each box plot. For the background, we conducted peak annotation over the entire CpG sites of each bulk.

### Supplementary Reference

[1] Chongyuan Luo, Christopher L Keown, Laurie Kurihara, Jingtian Zhou, Yupeng He, Junhao Li, Rosa Castanon, Jacinta Lucero, Joseph R Nery, Justin P Sandoval, et al. Single-cell methylomes identify neuronal subtypes and regulatory elements in mammalian cortex. Science, 357(6351):600–604, 2017.
